# A general model for analysis of linear and hyperbolic enzyme inhibition mechanisms

**DOI:** 10.1101/2025.04.10.648185

**Authors:** Rafael S. Chagas, Sandro R. Marana

## Abstract

The mechanisms of reversible inhibitors with a single binding site on enzymes are usually divided into basic groups: linear and hyperbolic, also called partial. Each of them subdivided into types: competitive, non-competitive and mixed. These six mechanisms are often considered separate identities. Here, prompted by the characterization of the inhibition of the wild-type and mutant β-glucosidase Sfβgly by Imidazole and Tris (2-amino-2-(hydroxymethyl)-1,3-propanediol) we developed a unifying enzyme kinetic model that integrates these six basic inhibition mechanism onto a single one. From this model we deduced a general enzyme kinetic equation that through modulation of simple parameters, i.e. the relative inhibitor affinity for two binding sites and the reactivity of the enzyme-substrate-inhibitor complex, is converted into the particular kinetic equation of each of those six inhibition mechanism. In short, we conclude that six fundamental inhibition mechanisms, linear and hyperbolic, are not isolate compartments, but actually facets of the same general model here presented.

## 1. Introduction

The mechanisms of the reversible inhibitors that have only a single binding site on enzymes are usually divided into two groups, linear and hyperbolic or partial. Both groups are called simple mechanisms due to the single binding site. Assuming initial rate (v_0_) conditions, they are delimited based on the behavior of the slope (*K*_s_/*k*_cat app_) and intercept (1/*k*_cat app_) lines in the Lineweaver-Burk plot (1/*v*_0_ x 1/[S]), which were determined under different inhibitor concentrations. So, as their denomination indicates, simple linear inhibitors render linear correlations when slope or intercept are plotted *versus* [I]. In agreement, simple hyperbolic (partial) inhibitors generate curved lines (hyperboles) in that plot, where the slope or intercept levels up at high [I].

These two groups can be further divided into types: competitive, non-competitive and mixed, which are identified by the relative positioning of the lines in the Lineweaver-Burk plots obtained with several [I]. Basically, these inhibition types differ about the positioning of the point in where the different lines cross. Hence, regardless the group, linear or hyperbolic, they are called intercepting mechanism.

Conversely, uncompetitive inhibitors are not intercepting because they produce parallel lines in the Lineweaver-Burk plots. Hence they belong to a different group.

In short, considering only the intercepting cases, there are six basic simple inhibition mechanisms, which are commonly represented in textbooks [1], but for clarity purpose they are also shown in the Supplementary Figure 1. Finally, as gathered from the classificatory effort, the inhibition mechanisms are understood as separate identities.

Initially, in a context of few structural data available, the inhibition mechanism studies provided a path to idealize schematic models of the enzyme structures. Nowadays, in a richer structural data environment, these inhibition models can be used to deposit additional layers of functional information on the enzymes structure. However, as these inhibition mechanistics models were not developed to represent microscopic molecular steps, as they are combined with structural data, tensions arise. Nevertheless, instead to represent a limitation, we saw these frictions as an opportunity to develop models amenable to smoothly integrate inhibition and structural binding data, harnessing the ease of the initial rate experiments to improve the functional characterization of enzymes.

Our starting point is the reversible inhibition of the GH1 β-glucosidase Sfβgly (PDB ID 5CG0; [2]) by Imidazole and Tris (2-amino-2-(hydroxymethyl)-1,3-propanediol) [3; 4]. Sfβgly, a digestive GH1 β-glucosidase from the fall armyworm *Spodoptera frugiperda*, has a tunnel-shaped active site, where short oligocellosaccharides (polymerization degree from 2 to 4), non-branched alkyl β-glucosides (4 to 12 carbons) and bulky aryl β-glucosides can fit [5]. The monosaccharide at the non-reducing extremity of the substrate (glycone) binds in the dead end of the tunnel (subsite -1), whereas the group (saccharide or not) linked to the glycone via the β-glycosidic bond is positioned towards the active site opening, a region that comprises the subsite +1 (Figure 1).

**Figure 1.**
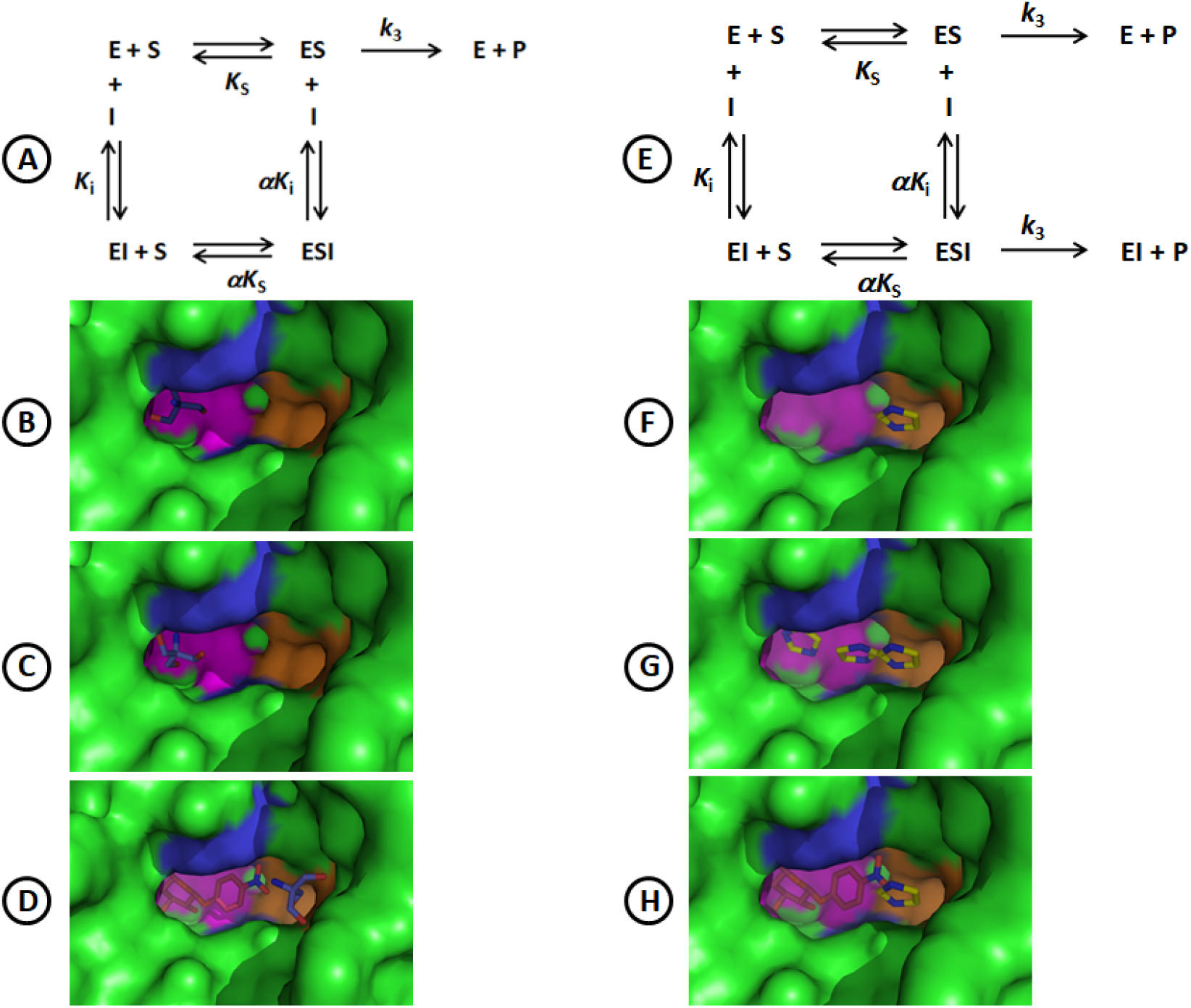
Mechanistic and structural description of the inhibition of the Sfβgly by Tris and Imidazole. Structures show a close up of the Sfβgly active site (PDB 5CG0). The subsite -1 is colored in pink, the subsite +1 in blue and a pocket in the lateral of the active site opening is marked in orange. **A)** Inhibition mechanism observed for Tris [4]. This mechanism is classified as intercepting, linear and mixed [1]. **B)** Tris docked within the subsite -1 of the Sfβgly active site. Based on [4]. **C)** Crystallographic structure for Tris bound within the Sfβgly active site (PDB 5CG0). Based on [3]. **D)** Putative ESI complex generated by docking of Tris (dark blue and red) in the Sfβgly active site previously occupied by NPβglc (p-nitrophenyl β-glucoside) (red). Tris is seen in the region of the pocket in the lateral of the active site opening (orange) **E)** Inhibition mechanism observed for Imidazole [3]. This mechanism is classified as intercepting, hyperbolic and partial competitive. F) Computational docking showing the Imidazole bound in a pocket in the lateral of the active site opening. Based on [3]. **G)** Simultaneous representation of three different docking solutions for Imidazole within the Sfβgly active site: subsite -1, subsite +1 and the pocket in the active site opening. Based on [3]. **H)** Putative ESI complex involving NPβglc (red), within the Sfβgly active site (subsites -1 and +1, pink and blue, respectively), and Imidazole (yellow and blue), in the pocket in the lateral of the active site opening (orange).

Imidazole acts as a simple hyperbolic competitive inhibitor, also known as partial competitive, while Tris functions as a simple linear mixed type (Figure 1). Despite their particularities and distinct identities, there is indeed some common ground between these mechanisms. For instance, both mechanisms present a ternary complex ESI, indicating that the inhibitor (either Imidazole or Tris) has a single binding site that is different from the substrate, which is found in both the free enzyme (E) and the enzyme-substrate complex (ES). Thus, this same single site is occupied by a single inhibitor molecule in the EI and ESI complexes. The same inhibition mechanism was observed when using synthetic and natural substrates, as p-nitrophenyl β-glucoside (NPβglc) and cellobiose, respectively [3; 4].

In agreement, computational docking suggested that a pocket on the side of the Sfβgly active site opening could be the binding site for Imidazole, leaving the -1 and +1 subsites available for the substrate (Figure 1). Hence, such lateral binding site would be compatible with the formation of the ESI complex, which, as noted above, is a requirement for the partial competitive inhibition mechanism observed experimentally [3; 1] (Figure 1). The crater-shaped opening of the Sfβgly active site, adapted to bind bulky aryl groups as seen in the glycoside amygdalin and methylumbelliferyl β-glucoside [6], is indeed large enough to comprise that pocket in its lateral and to enable the simultaneous interaction with the substrate and the inhibitor, as proposed in the putative ESI complex involving NPβglc and Imidazole suggested by computational docking (Figure 1). Finally, residues S247, N249 and F251, placed in that pocket, interact exclusively with Imidazole (Figure 1) [3].

Nevertheless, computational docking data pointed that Imidazole could also potentially bind at several sites along the Sfβgly active site, from the -1 subsite, at the dead end of the active site, up to +1 subsite, close to its wide opening, in addition to the lateral pocket [3] (Figure 1).

Tris is bound within the -1 subsite of the Sfβgly crystallographic structure [2], an interaction also detected by computational docking. However, Tris can also bind to a site in the vicinity of the lateral pocket in the active site opening when docked with the NPβglc-Sfβgly complex (Figure 1). So, whereas the binding within the -1 subsite is typical of a competitive inhibition, which is not observed experimentally for Tris and Sfβgly, the interaction around the active site opening is consistent with the detection of a linear mixed inhibition mechanism, which includes the formation of an unproductive ESI complex.

Nevertheless, a picture combining these observations results in some tension, which is detailed in the next lines. The interactions of Imidazole and Tris around the Sfβgly active site opening are actually compatible with their experimental inhibition mechanisms, which include the formation of ESI complexes (Figure 1). As a consequence, this consideration would imply that the interactions of Imidazole and Tris in the internal portion of the active site (−1 and +1 subsites) should be irrelevant, because they do not influence the inhibition mechanism. But, the experimental detection of Tris within the -1 subsite of the Sfβgly crystallographic structure suggests otherwise. This apparent contradiction could be solved if Imidazole and Tris could really interact with multiple spots along the active site, as shown by their computational dockings (Figure 1). However, this Imidazole or Tris mode of binding at multiple sites is apparently incompatible with their experimental inhibition mechanisms, which are based on the interaction of each inhibitor at a single site (Figure 1).

To dissipate such tension from the picture that combines the binding of Imidazole and Tris to Sfβgly with their observed inhibition mechanisms, we followed an experimental and theoretical approach.

Single mutations were introduced at Sfβgly residues S247, N249 and F251, aiming to alter Imidazole interaction within the lateral pocket in the active site opening. Residue S247 was replaced by Y, N249 was exchanged by Q and A and F251 was swept by A. The inhibitory effect of Imidazole and Tris on the mutant Sfβgly was characterized and compared to the wild-type enzyme.

Finally, to interpret our experimental observations, we developed a general inhibition model that unifies six fundamental inhibition mechanisms (linear and hyperbolic) previously described in the literature. This general model not only explained the inhibitory effect of Imidazole and Tris on Sfβgly, but it is also an effective tool for characterizing the mechanisms of a broad range of small molecule inhibitors presenting single or multiple binding sites in enzymes.

## 3. Results

The Sfβgly mutants were produced as recombinant proteins in *E. coli* and purified following the procedures previously published (Supplementary Figure 2) [3]. These enzyme samples were used to determine the initial rate of hydrolysis of different concentrations of NPβglc in the presence of either Imidazole or Tris, each tested separately at several concentrations. The data were analyzed using the Lineweaver-Burk plot to determine the inhibition mechanism, a classical enzyme kinetics approach [1].

Next, the effects of each mutation on the inhibitors will be presented separately. The findings will then be combined and discussed.

### Mutational effects on Imidazole inhibition

The mutations in the residues that form the lateral pocket at the active site opening produced diverse effects on Imidazole inhibition mechanisms.

The partial competitive mechanism, originally observed for the wild-type Sfβgly, changed to a linear mixed mechanism for the S247Y mutant (Figure 2). This change is indicated by the lines crossing in the second quadrant of the Lineweaver-Burk plot, as well as by the linear effect of the [I] on the *K*_s_/*k*_cat app_, derived from the lines slope, and also on the 1/*k*_cat app_, derived from the lines intercept in the 1/ ν_0_ axis [1].

**Figure 2.**
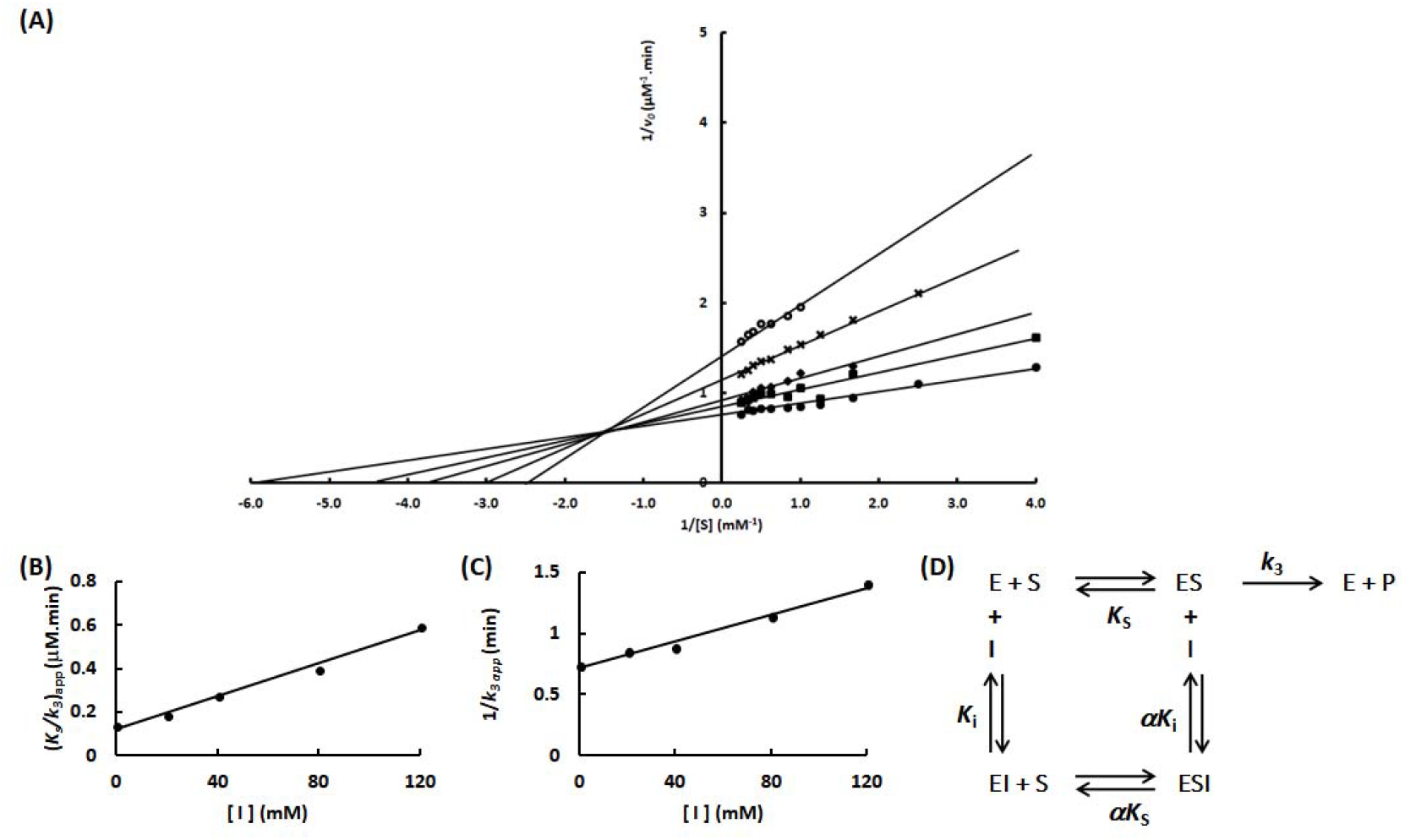
Determination of the Imidazole inhibition mechanism in the Sfβgly S247Y mutant. **(A)** Effect of different Imidazole concentrations (⍰, 0; ▪, 20 mM; ♦, 40 mM; x, 80 mM; ⍰, 120 mM) on the initial rate of hydrolysis of the substrate NPβglc shown in Lineweaver-Burk plots. **(B)** Effect of the Imidazole concentration on the apparent K_s_/k_3_ (calculated from the line slope). **(C)** Effect of the Imidazole concentration on the apparent 1/ k_3_ (calculated from the line intercept). Rates are the mean of three determinations of the product formed using the same enzyme sample. **(D)** – Linear mixed inhibition mechanism (intersecting, linear, mixed). S, substrate NPβglc; I, inhibitor Imidazole; E, enzyme Sfβgly; P, product; *K*_s_, dissociation constant for the ES complex; *K*_i_, dissociation constant for the EI complex; α*K*_i_, dissociation constant for the ESI complex; α, factor that represents the mutual hindering effect between S and I (α >1); *k*_3_, rate constant for product formation [1].

Besides that, a linear competitive mechanism was observed for the N249Q and N249A mutants (Figures 3 and 4). As shown, the Lineweaver-Burk plots for these mutants show a set of lines with escalating slope as the inhibitor concentration increases, while those lines converge to the same intercept at the 1/*ν*_0_ axis regardless of the inhibitor concentration. Additionally, plots of the apparent *K*_s_/*k*_cat_ (calculated from the lines slopes) versus the inhibitor concentrations are linear (Figures 3 and 4). These are typical features of a linear competitive inhibition mechanism [1].

**Figure 3.**
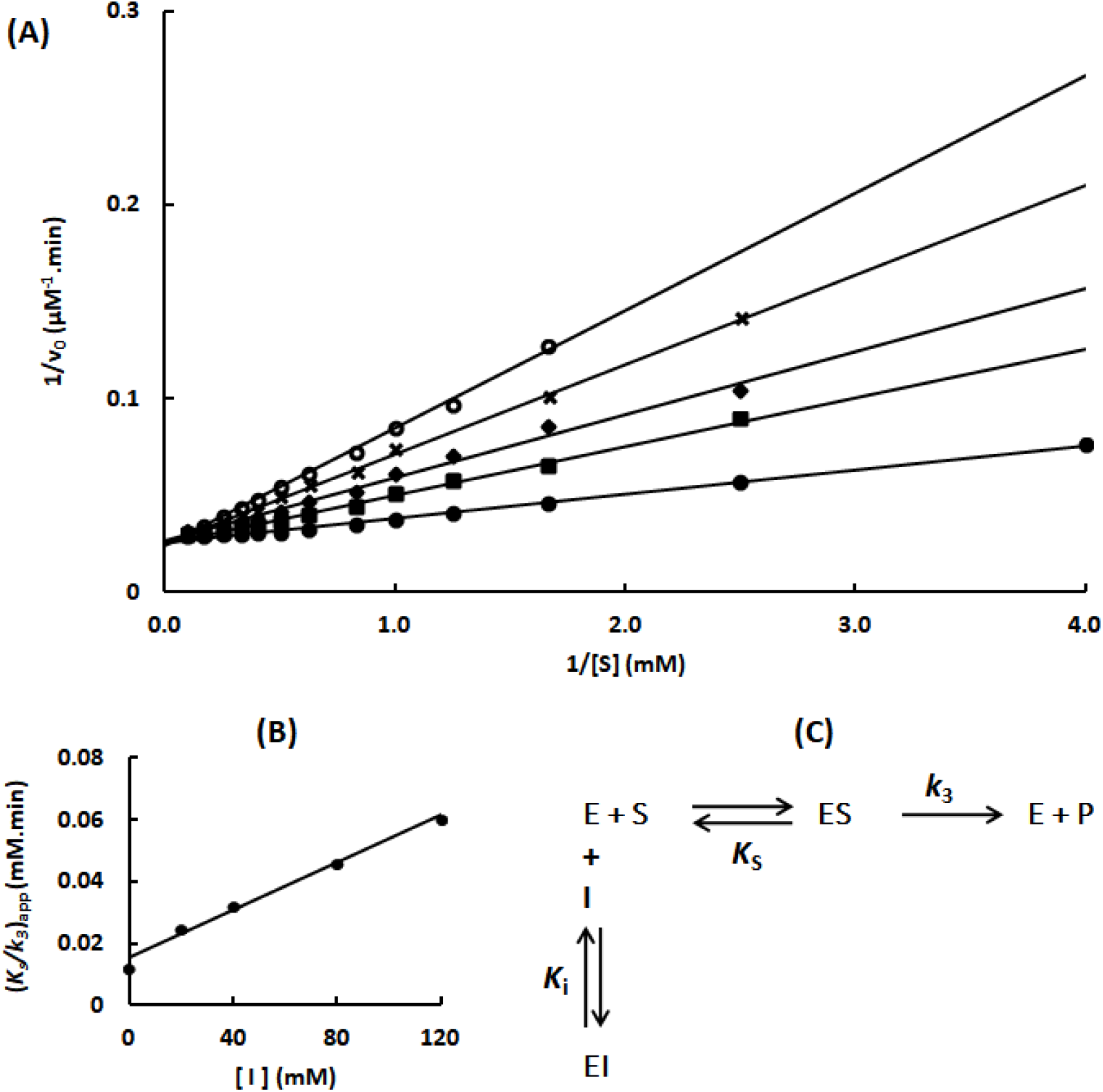
Determination of the Imidazole inhibition mechanism in the Sfβgly N249Q mutant. **(A)** Effect of different Imidazole concentrations ⍰, 0; ▪, 20 mM; ♦, 40 mM; x, 80 mM; ⍰, 120 mM) on the initial rate of hydrolysis of the substrate NPβglc shown in Lineweaver-Burk plots. **(B)** Effect of the Imidazole concentration on the apparent *K*_s_/*k*_3_ (calculated from the line slope). Rates are the mean of three determinations of the product formed using the same enzyme sample. **(C)** Linear competitive inhibition mechanism (intersecting, linear, competitive). S, substrate NPβglc; I, inhibitor Imidazole; E, enzyme Sfβgly; P, product; *K*_s_, dissociation constant for the ES complex; *K*_i_, dissociation constant for the EI complex; *k*_3_, rate constant for product formation [1].

**Figure 4.**
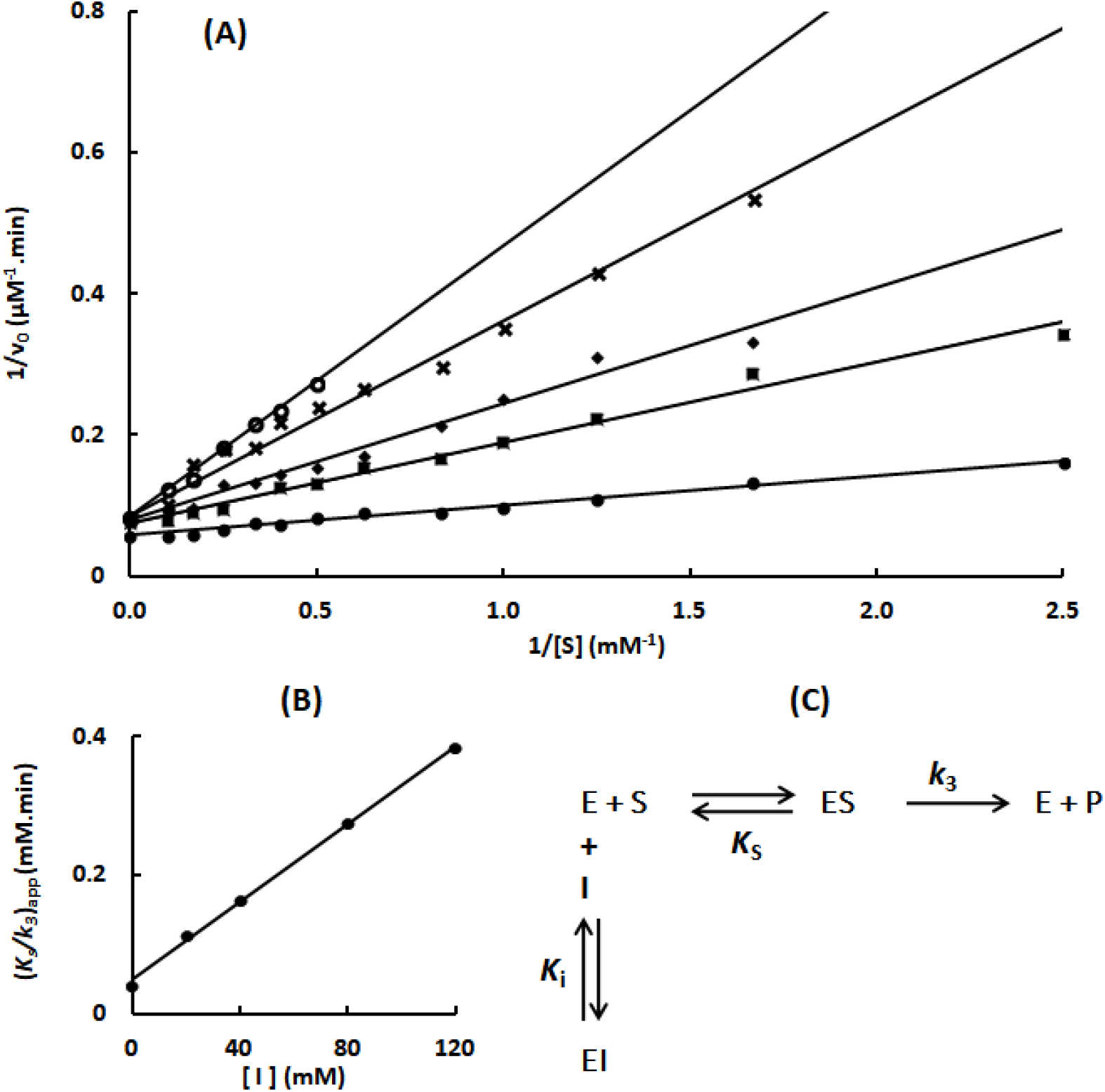
Determination of the Imidazole inhibition mechanism in the Sfβgly N249A mutant. **(A)** Effect of different Imidazole concentrations ⍰, 0; ▪, 20 mM; ♦, 40 mM; x, 80 mM; ⍰, 120 mM) on the initial rate of hydrolysis of the substrate NPβglc shown in Lineweaver-Burk plots. **(B)** Effect of the Imidazole concentration on the apparent *K*_s_/*k*_3_ (calculated from the line slope). Rates are the mean of three product determination using the same enzyme sample.**(C)** Linear competitive inhibition mechanism (intersecting, linear, competitive). S, substrate NPβglc; I, inhibitor Imidazole; E, enzyme Sfβgly; P, product; *K*_s_, dissociation constant for the ES complex; *K*_i_, dissociation constant for the EI complex; *k*_3_, rate constant for product formation [1].

Finally, a partial mixed mechanism was observed for the F251A mutant (Figure 5), which is characterized by lines crossing in the second quadrant of the Lineweaver-Burk plot and hyperbolic plots of *K*_s_/*k*_cat app_ versus [I] and 1/*k*_cat app_ versus [I]. In addition, these hyperbolic plots can be converted into lines by calculating the variation of the slope (Δ *K*_s_/*k*_cat app_) and intercept (Δ 1/*k*_cat app_) due to [I] increment taking the lines in absence of inhibitor as reference.

**Figure 5.**
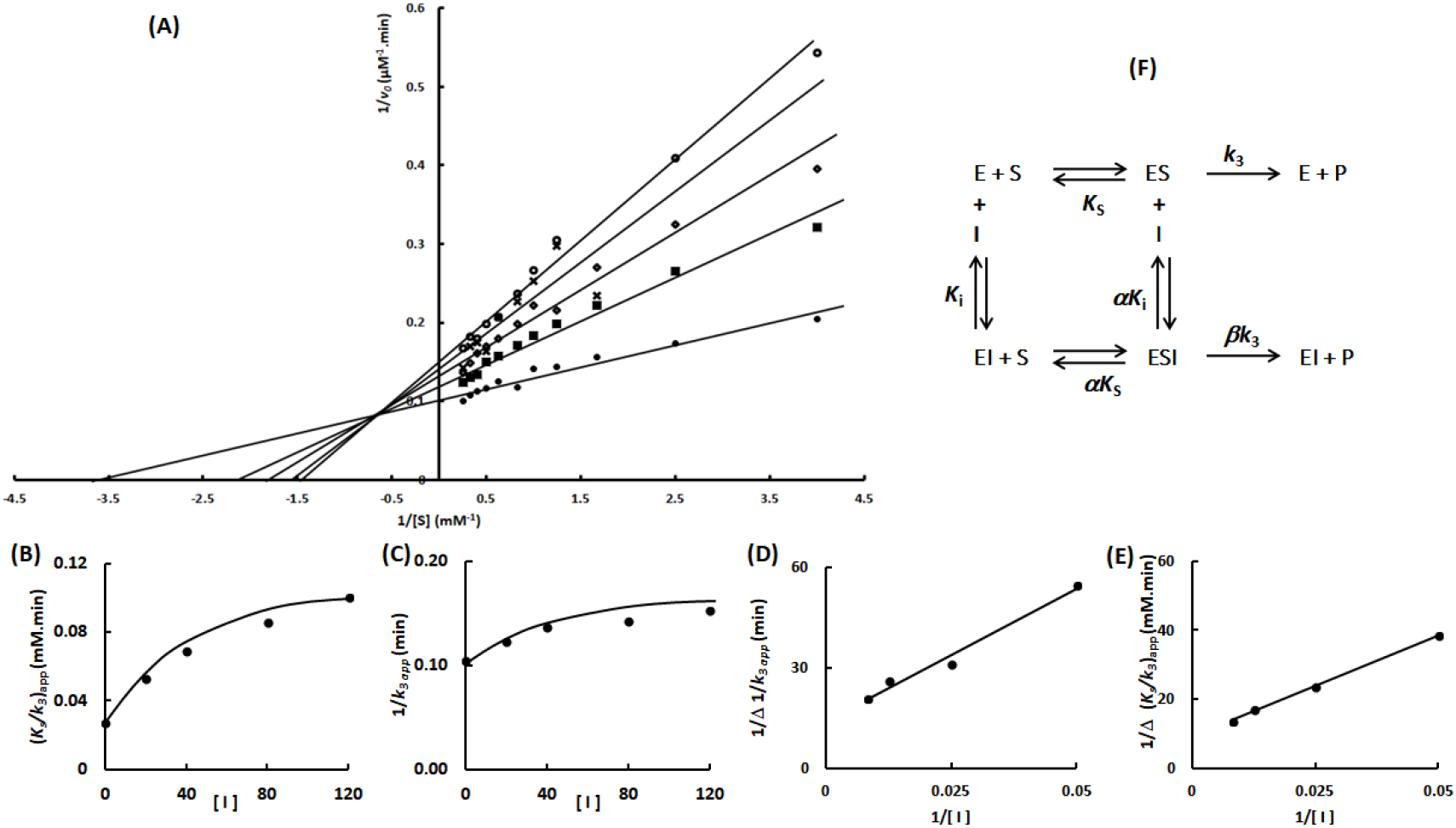
Determination of the Imidazole inhibition mechanism in the Sfβgly F251A mutant. **(A)** Effect of different Imidazole concentrations ⍰, 0; ▪, 20 mM; ♦, 40 mM; x, 80 mM; ⍰, 120 mM) on the initial rate of hydrolysis of the substrate NPβglc shown in Lineweaver-Burk plots. **(B)** Effect of the Imidazole concentration on the apparent *K*_s_/*k*_3_ (calculated from the line slope). (C) Effect of the Imidazole concentration on the apparent 1/*k*_3_ (calculated from the line intercept). (D) Secondary plot of 1/Δ_*K*s/*k*3_ *versus* 1/[ I]. Δ_*K*s/*k*3_ = app *K*_s_/*k*_3 i_ – app *K*_s_/*k*_3 0_, in which app *K*_s_/*k*_3 i_ corresponds to the line in the presence of imidazole and app *K*_s_/*k*_3 0_ to the line in the absence of Imidazole. **(E)** Secondary plot of 1/Δ_1/ *k*3_ versus 1/[ I]. Δ_1/ *k*3_ = app 1/*k*_3 i_ – app 1/*k*_3 0_, in which app 1/*k*_3 i_ corresponds to the line in the presence of imidazole and app 1/*k*_3 0_ to the line in the absence of Imidazole. Rates are the mean of three product determination using the same enzyme sample. (F) Hyperbolic mixed inhibition mechanism (intersecting, hyperbolic, mixed). S, substrate NPβglc; I, inhibitor Imidazole; E, enzyme Sfβgly; P, product; *K*_s_, dissociation constant for the ES complex; *K*_i_, dissociation constant for the EI complex; α*K*_i_, dissociation constant for the ESI complex; α factor represents the mutual hindering effect between S and I (α >1); *k*_3_, rate constant for product formation complex from the ES complex; β*k*_3_, rate constant for product formation from the ESI complex, (β <1) [1].

Suitable parameters for describing each Imidazole inhibition mechanism were calculated based on the plots of *K*_s_/*k*_cat app_ *versus* [I] and 1/*k*_cat app_ *versus* [I] as previously established [1] (Table 1).

**Table 1.**
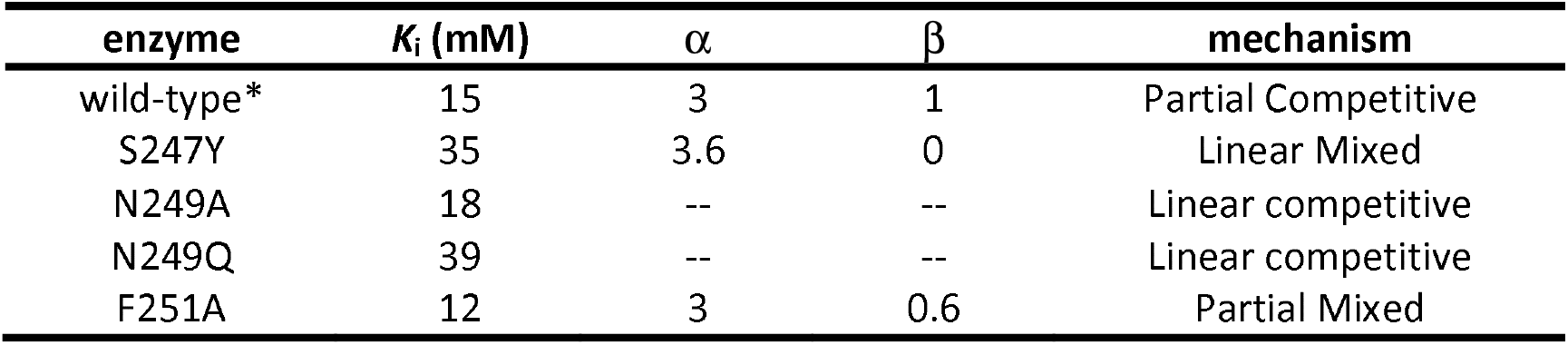
Parameters of the Imidazole inhibition mechanism for wild-type and Sfβgly mutants

| enzyme | $K_i$ (mM) | $\alpha$ | $\beta$ | mechanism |
| --- | --- | --- | --- | --- |
| wild-type* | 15 | 3 | 1 | Partial Competitive |
| S247Y | 35 | 3.6 | 0 | Linear Mixed |
| N249A | 18 | -- | -- | Linear competitive |
| N249Q | 39 | -- | -- | Linear competitive |
| F251A | 12 | 3 | 0.6 | Partial Mixed |

The inhibition mechanisms are depicted on Figures 2 and 5. *K*_i_ represents the dissociation constant of the Sfβgly enzyme-Imidazole complex (EI). α corresponds to the mutual hindering effect involving substrate and Imidazole. α does not apply for the simple competitive mechanism. β represents the Imidazole effect on the *k*_3_. β < 1 indicates that the ESI complex is less productive than the ES complex. β = 1 shows that ES and ESI are equally productive. β = 0 indicates that the ESI complex is not productive [1]. *Data for the wild-type Sfβgly were taken from [3].

### Mutational effects on Tris inhibition

The mutational effects on the Tris inhibition mechanism were less diverse. Originally, Tris acted as a linear mixed inhibitor of the wild-type Sfβgly (Figure 1). However, S247Y, N249Q and N249A mutations converted it to a linear competitive inhibitor (Figures 6 to 8). Thus, *K*_i_ value for Tris with these mutants were calculated based on the *K*_s_/*k*_cat app_ *versus* [I] plots [1] (Table 2).

**Table 2.** Parameters of the Tris inhibition mechanism for wild-type and Sfβgly mutants

| enzyme | $K_i$ (mM) | $\alpha$ | $\beta$ | mechanism |
| --- | --- | --- | --- | --- |
| wild-type* | 12 | 3 | 0 | Linear Mixed |
| S247Y | 71 | -- | -- | Linear competitive |
| N249A | 0.6 | -- | -- | Linear competitive |
| N249Q | 32 | -- | -- | Linear competitive |
| F251A | 10 | 12 | 0 | Linear Mixed |

**Figure 6.**
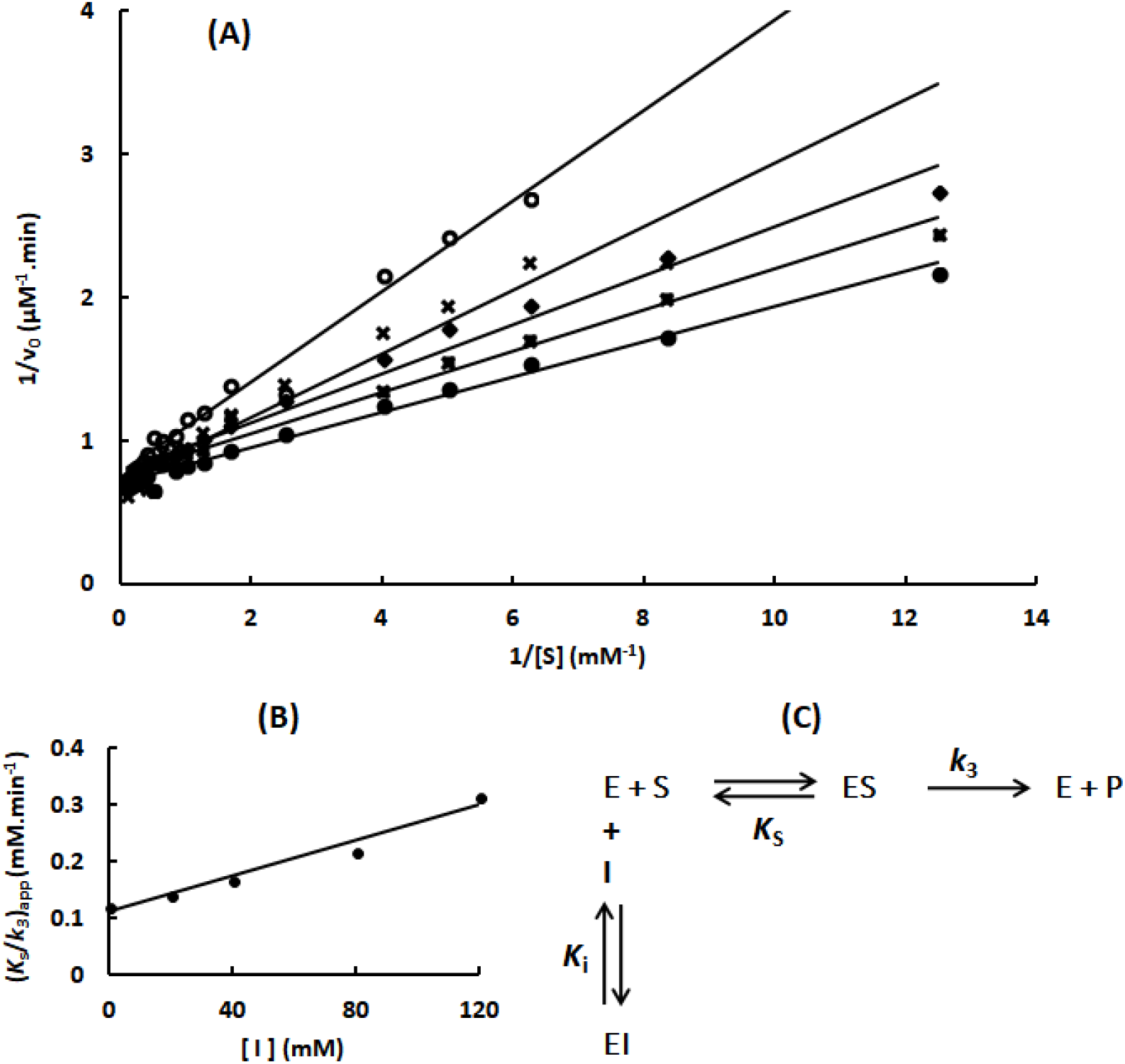
Determination of the Tris inhibition mechanism in the Sfβgly S247Y mutant. **(A)** Effect of different Tris concentrations ⍰, 0; ▪, 20 mM; ♦, 40 mM; x, 80 mM; ⍰, 120 mM) on the initial rate of hydrolysis of the substrate NPβglc shown in Lineweaver-Burk plots. **(B)** Effect of the Tris concentration on the apparent *K*_s_/*k*_3_ (calculated from the line slope). Rates are the mean of three product determination using the same enzyme sample. (C) Linear competitive inhibition mechanism (intersecting, linear, competitive). S, substrate NPβglc; I, inhibitor Tris; E, enzyme Sfβgly; P, product; *K*_s_, dissociation constant for the ES complex; *K*_i_, dissociation constant for the EI complex; *k*_3_, rate constant for product formation [1].

Conversely, the F251A mutation had no effect on the inhibition mechanism for Tris, which remained a linear mixed type (Figure 8). Accordingly, plots of *K*_s_/*k*_cat app_ *versus* [I] and 1/*k*_cat app_ *versus* [I] were used to determine the *K*_i_ and the α factor values for Tris in the F251A mutant [1] (Table 1).

The inhibition mechanisms are depicted on Figures 6 and 8. *K*_i_ represents the dissociation constant of the Sfβgly enzyme-Tris complex (EI). α corresponds to the mutual hindering effect involving substrate and Tris. α does not apply for the simple competitive mechanism. β represents the Tris effect on the *k*_cat_. β = 0 indicates that the ESI complex is not productive [1]. *Data for the wild-type Sfβgly were taken from [4].

## Discussion

### Mutations lead to the emergence of different inhibition mechanisms

Tables 1 and 2 show a remarkable alteration of the inhibition mechanism for both, Imidazole and Tris, due to mutations in the lateral pocket of the active site. For instance, all 4 mutations here analyzed changed the Imidazole inhibition mechanism resulting in the emergence of three distinct mechanisms (Table 1). Besides that, 3 out of 4 mutations converted Tris into a linear competitive inhibitor of the Sfβgly (Table 2). Conversely, clear modifications in K_i_ are few. For example, five-fold changes were only observed for the S247Y and N249A mutants when using Tris as inhibitor (Table 2). Besides that, S247Y and N249Q mutations resulted in a two-fold increase in the *K*_i_ for Imidazole and Tris, respectively (Table 1 and 2). However, given that the inhibition mechanism changed in these cases, it is not unambiguous that *K*_i_ would be describing the inhibitor interaction within the same site in the wild-type and mutant Sfβgly.

In short, contrary to eventual expectations, the effects of the replacement in the residues that form the lateral pocket of the active site were not mainly restricted to inhibitor affinity changes. Actually, those replacements allowed different inhibition mechanism to emerge from those originally observed for the wild-type Sfβgly. This remarkable finding has to be carefully analyzed.

The imidazole inhibition mechanism, which is partial competitive for the wild-type Sfβgly, changed to linear competitive in the N249A and N249Q mutants (Table 1; Figures 1, 3 and 4). The partial competitive mechanism involves the presence of both EI and ESI complexes (Figure 1). Hence, in the wild-type Sfβgly, Imidazole binds to a single and same site present in both E and ES, which is compatible to the proposal that such site is the lateral pocket in the opening of the active site [3]. Moreover, imidazole binding at this site does not obstruct substrate binding in the inner region of the active site. On the other hand, the linear competitive mechanism observed for the N249A and N249Q mutants indicates the occurrence of an EI complex, where substrate binding is impeded (Figures 3 and 4). Hence, the EI complexes present in these two mechanisms, despite being represented by the same symbol, are not physically equivalent. In the first mechanism, the Imidazole interaction site within the Sfβgly is distinct from the substrate binding site, whereas in the second mechanism, both the inhibitor and the substrate bind at the same spot. Therefore, a linear competitive inhibition mechanism is not embedded in a partial competitive one.

Tris was originally a linear mixed-type inhibitor of the wild-type Sfβgly (Figure 1). This mechanism means that Tris also finds a same single binding site in E and ES forming complexes EI and ESI. Conversely, the linear competitive inhibition observed with the Tris in S247Y, N249Q and N249A mutants indicates the presence of only an EI complex, where substrate binding is excluded (Table 2; Figures 6, 7 and 8). Hence, the EI complexes present in these two mechanisms are also not equivalent. In short, a linear competitive inhibition mechanism is not contained in a linear mixed one.

**Figure 7.**
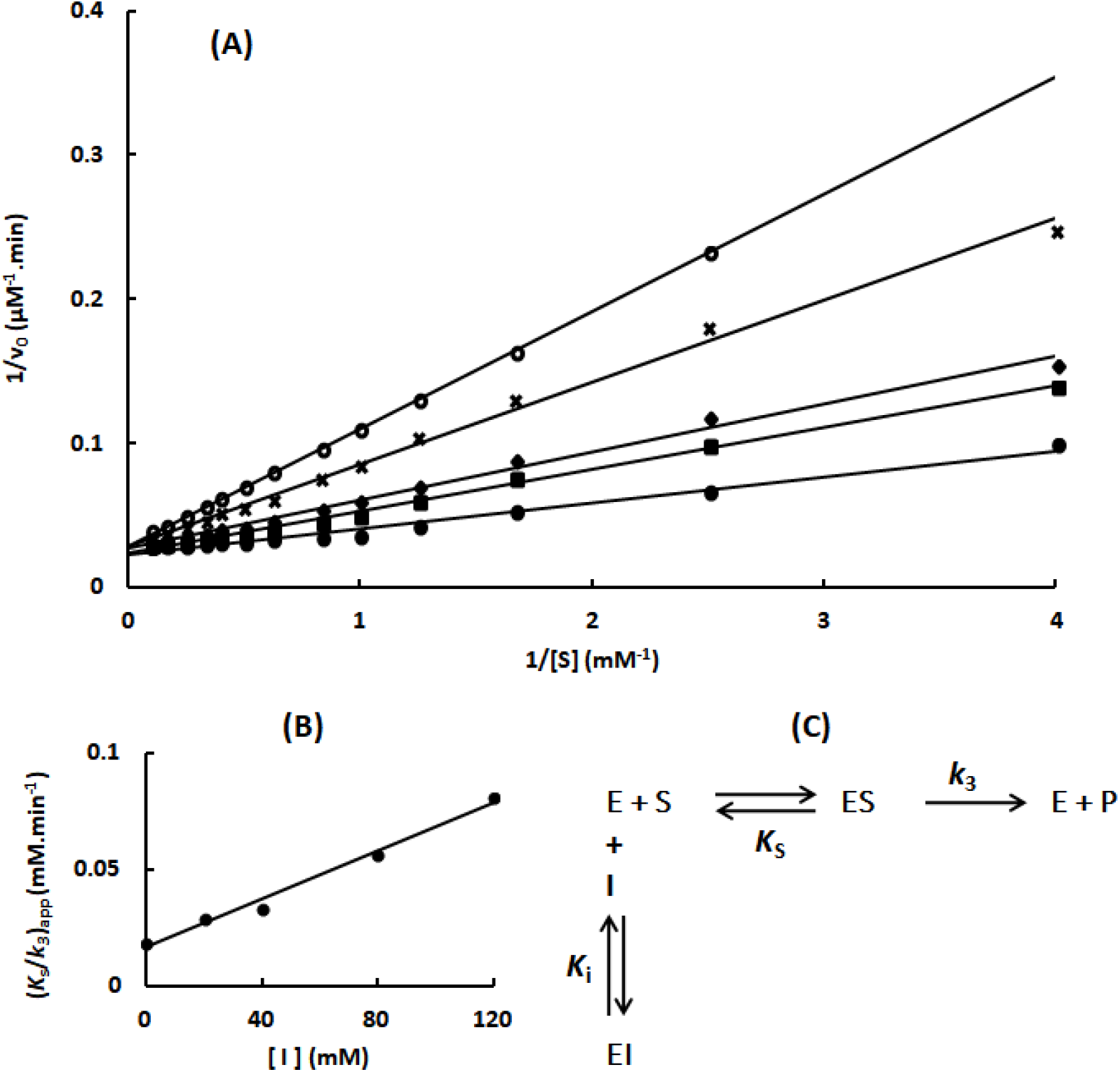
Determination of the Tris inhibition mechanism in the Sfβgly N249Q mutant. **(A)** Effect of different Tris concentrations (⍰, 0; ▪, 20 mM; ♦, 40 mM; x, 80 mM; ⍰, 120 mM) on the initial rate of hydrolysis of the substrate NPβglc shown in Lineweaver-Burk plots. **(B)** Effect of the Tris concentration on the apparent *K*_s_/*k*_3_ (calculated from the line slope). Rates are the mean of three product determination using the same enzyme sample. **(C)** Linear competitive inhibition mechanism (intersecting, linear, competitive). S, substrate NPβglc; I, inhibitor Tris; E, enzyme Sfβgly; P, product; *K*_s_, dissociation constant for the ES complex; *K*_i_, dissociation constant for the EI complex; k_3_, rate constant for product formation [1].

**Figure 8.**
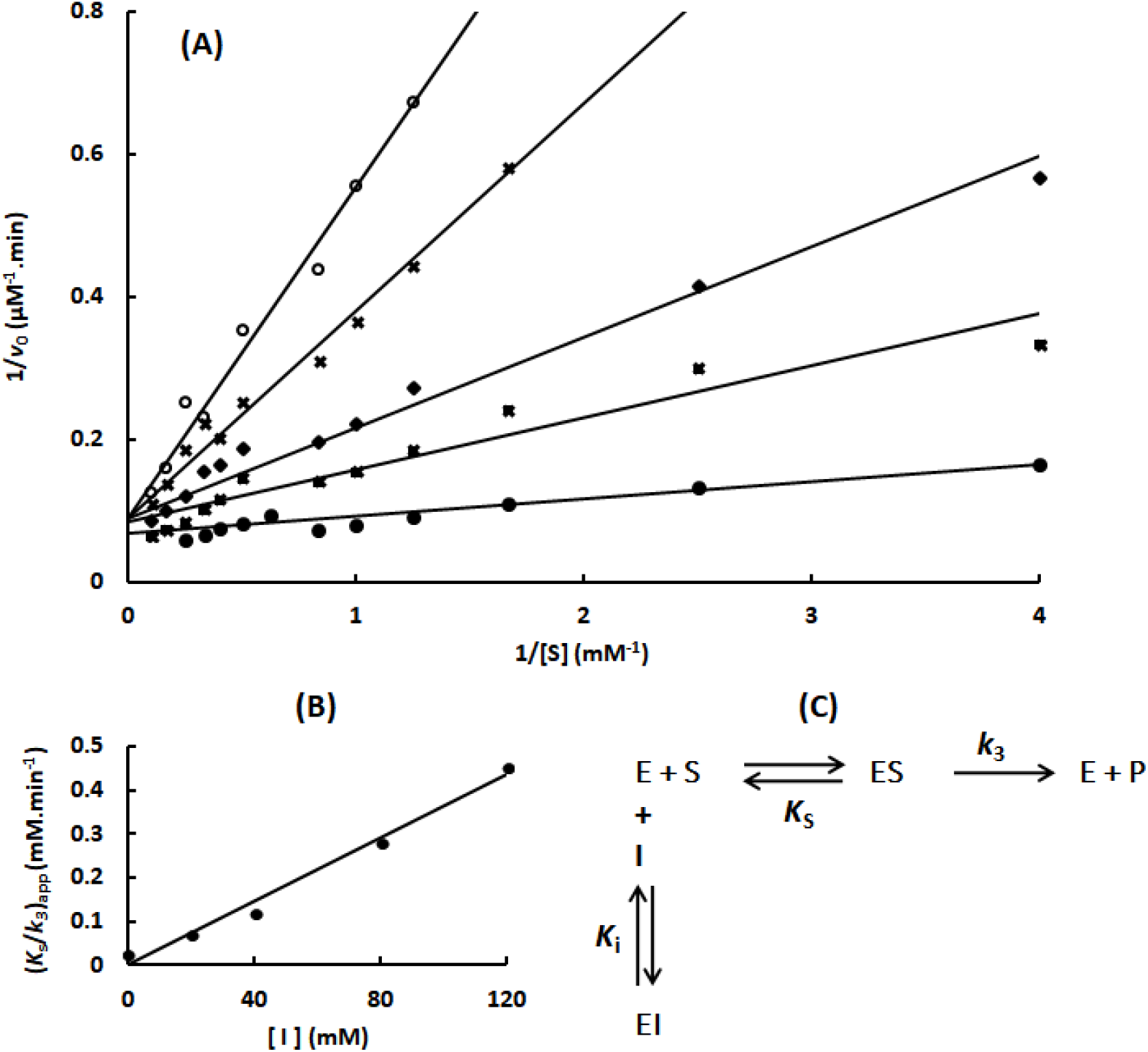
Determination of the Tris inhibition mechanism in the Sfβgly N249A mutant. **(A)** Effect of different Tris concentrations (⍰, 0; ▪, 20 mM; ♦, 40 mM; x, 80 mM; ⍰, 120 mM) on the initial rate of hydrolysis of the substrate NPβglc shown in Lineweaver-Burk plots. **(B)** Effect of the Tris concentration on the apparent *K*_s_/*k*_3_ (calculated from the line slope). **(C)** Effect of the Tris concentration on the apparent 1/ *k*_3_ (calculated from the line intercept). Rates are the mean of three product determination using the same enzyme sample. **(D)** Linear competitive inhibition mechanism (intersecting, linear, competitive). S, substrate NPβglc; I, inhibitor Tris; E, enzyme Sfβgly; P, product; *K*_s_, dissociation constant for the ES complex; *K*_i_, dissociation constant for the EI complex; *k*_3_, rate constant for product formation [1].

**Figure 9.**
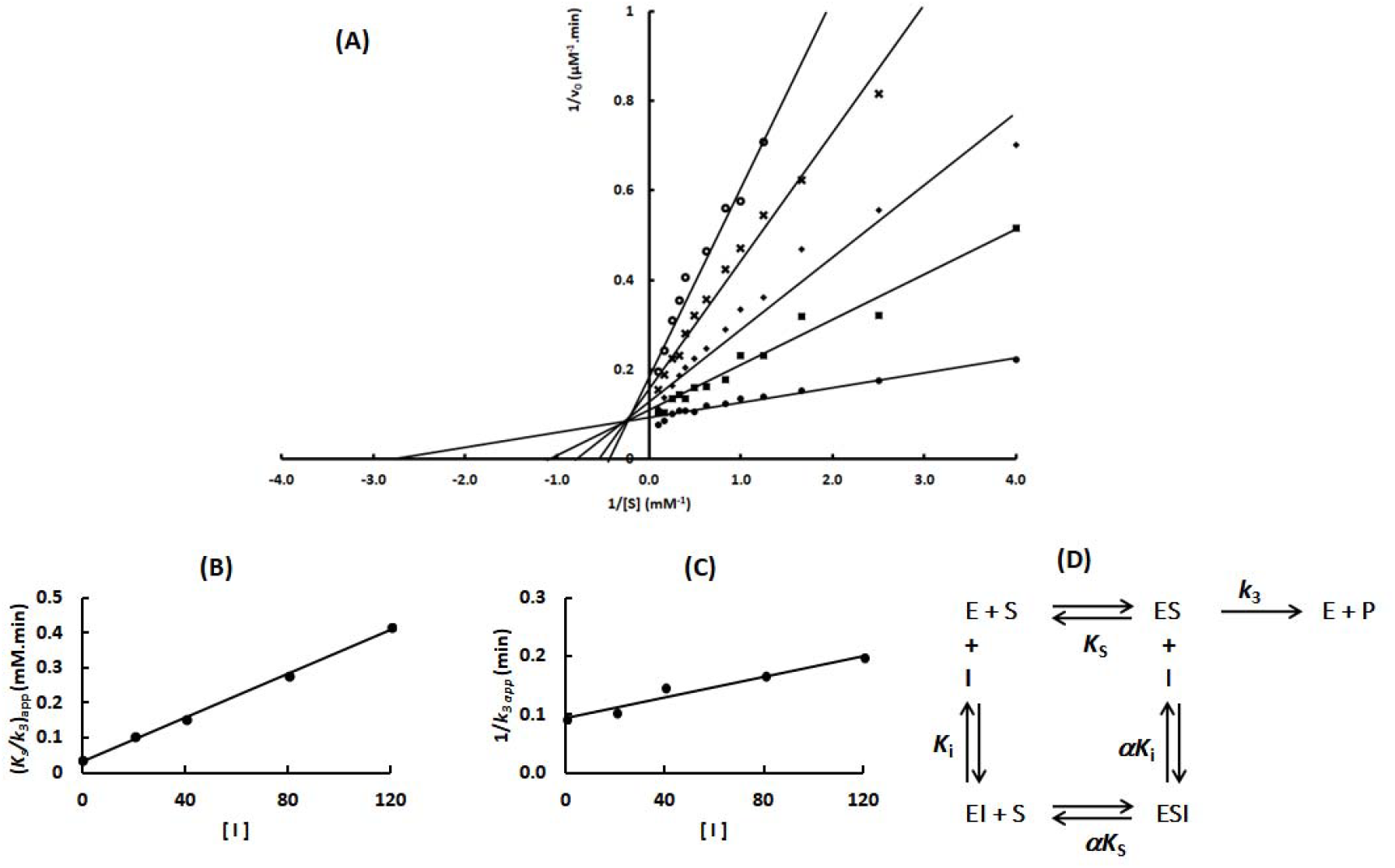
Determination of the Tris inhibition mechanism in the Sfβgly F251A mutant. **(A)** Effect of different Tris concentrations (⍰, 0; ▪, 20 mM; ♦, 40 mM; x, 80 mM; ⍰, 120 mM) on the initial rate of hydrolysis of the substrate NPβglc shown in Lineweaver-Burk plots. **(B)** Effect of the Tris concentration on the apparent *K*_s_/*k*_3_ (calculated from the line slope). (C) Effect of the Tris concentration on the apparent 1/ *k*_3_ (calculated from the line intercept). Rates are the mean of three product determination using the same enzyme sample. (D) Linear mixed inhibition mechanism (intersecting, linear, mixed). S, substrate NPβglc; I, inhibitor Tris; E, enzyme Sfβgly; P, product; *K*_s_, dissociation constant for the ES complex; *K*_i_, dissociation constant for the EI complex; α*K*_i_, dissociation constant for the ESI complex; α, factor that represents the mutual hindering effect between S and I (α >1); *k*_3_, rate constant for product formation [1].

Therefore, considering that the inhibition mechanisms that emerge for the Sfβgly mutants mentioned above (Table 1 and 2) were not already contained in those observed for the wild-type enzyme, they could not merely result from total or partial mutational damage in the original binding site of the inhibitor. In fact, those emerging mechanism demand a different binding site for the inhibitor. Nevertheless, if that different binding site were previously in place in the wild-type Sfβgly, it also should be occupied by the inhibitor, which in turn is not compatible to the single binding site inhibition mechanism experimentally detected for Imidazole and Tris (Table 1 and 2).

### Development of a general inhibition mechanism

Hence, the emergence of unexpected inhibition mechanism for the Sfβgly mutants is pointing that the mechanisms observed for the wild-type enzyme have to be interpreted from a new perspective. Based on this, we propose here that these results could be rationalized by assuming that the inhibition mechanism observed for the wild-type Sfβgly in the presence of Imidazole or Tris actually arise from the combination of the same inhibitor interaction with at least two different sites.

Though, such combination has to adjust to the following criteria. The two different mechanisms, each one reflecting the inhibitor interaction in a different site, have to blend mimicking a recognizable inhibition mechanism previously observed for the wild-type enzyme and characterized by the inhibitor interaction within a single site. Then, these two putative separate mechanisms must merge into a single kinetic equation that describes the same behavior of the lines in the Lineweaver-Burk plot previously observed for the wild-type Sfβgly. Finally, such combination should be compatible with the emergence of different inhibition mechanisms for both Imidazole and Tris upon mutation of the wild-type Sfβgly.

The large number of initial possibilities to be combined is passable to reduction by observing the outcomes of the molecular docking for Imidazole and Tris in the Sfβgly structure. Basically, Imidazole and Tris are observed interacting in three different regions: the -1 subsite, at the dead end of the active site; the +1 subsite, at the active site opening to the protein surface and the pocket located laterally to the active site opening [3; 4] (Figure 1). The inhibitor interactions within -1 and +1 subsites should obstruct the substrate binding, so from that perspective they can be seen as a single binding site. On the other hand, interactions near to the active site opening and in its lateral pocket could impair the product formation or even to be innocuous in that aspect, but it would not interfere in the substrate binding. Therefore, the simplest starting mechanism combines an inhibitor interaction within the active site, which hinders the substrate binding, and an additional interaction beyond that spot with no effect on the substrate interaction (Figure 10).

**Figure 10.**
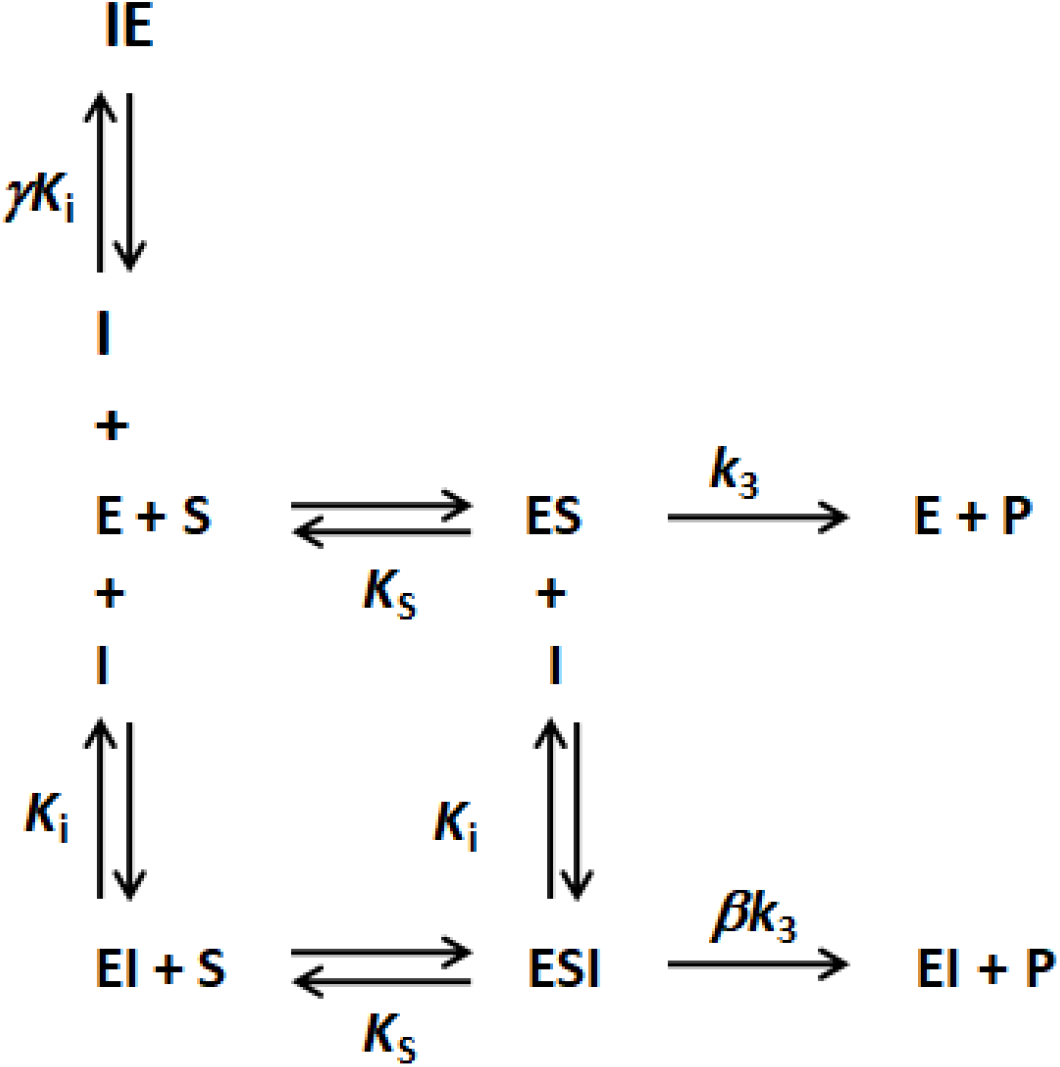
General inhibition mechanism considering two different binding sites for the inhibitor within the enzyme. One of the sites is within the active site, where the inhibitor interaction blocks the substrate binding and results in the IE complex. The second site is beyond the active site, where the inhibitor binding does not affect the substrate interaction within the active site. This binding produces the EI complex. E, free enzyme; ES, enzyme-substrate complex; IE, complex presenting the inhibitor bound within the enzyme active site; EI, complex in which the inhibitor is interacting beyond the active site; ESI, ternary complex presenting the substrate in the enzyme active site and the inhibitor bound elsewhere; γ factor represents the inhibitor relative affinity for the two binding sites available in the enzyme; β represents the inhibitor effect on the *k*_3_.

Figure 10 depicts these basic premises of the putative interaction between Imidazole or Tris and Sfβgly. The inhibitor binding at the active site opening in the free E or ES complex produces the EI and ESI complexes; an interaction represented by the dissociation constant, *K*_i._ This inhibitor-enzyme interaction does not affect the substrate binding, so its affinity is the same for the free E and the complex EI, which is then expressed as *K*_s_. The relative reactivity of the ES and ESI complexes is expressed in the β factor. If both are equally productive, then β = 1 and *k*_3_ and β*k*_3_ are the same rate constant. Nevertheless, if ESI does not generate product, it follows that β = 0. Finally, the range 0 < β < 1 indicates that inhibitor hinders the product formation, so ESI is less productive than ES (βk_3_ < *k*_3_). In addition to that, Imidazole or Tris binding within the active site per se, the subsites -1 or +1, form the IE complex, an equilibrium mediated by the γ*K*_i_ dissociation constant. The factor γ multiplying *K*_i_ was introduced to express the relative affinity of the same inhibitor for these two different sites, i.e. the region around the active site opening and within the active site. Thus if γ = 1, the inhibitor presents the same affinity for both sites. Conversely γ < 1 indicates that the inhibitor binds preferentially in the the active site. An extreme value, γ << 1, points to an exclusive interaction within the active site. The opposite is true for γ factor higher than 1.

Based on these premises, a general kinetic equation expressing the effect of the [S] on the initial rate in the presence of a [I] was deduced (equation 1) (Supplementary Figure 3):

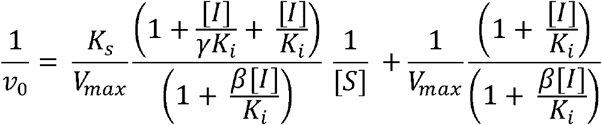

### The general model explains the Imidazole and Tris inhibition of the wild-type Sfβgly

To meet the criteria mentioned above, the general model (Figure 10 and Equation 1) has to produce data that could be interpreted as partial competitive and linear mixed mechanisms, which were observed for Imidazole and Tris when inhibiting the wild-type Sfβgly, respectively (Table 1 and 2)[3; 4].

Hence assuming a particular situation in which the inhibitor is totally innocuous regarding the product formation, β = 1, the equation 1 simplifies to equation 2 (Supplementary Figure 3):

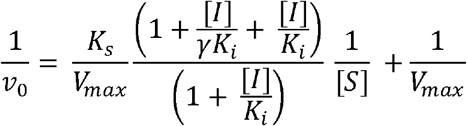

The equation 2 shows an intercept that does not depend on [I]. So, assuming equally productive ES and ESI, the general mechanism (Figure 10) results in lines that converge to the same point at the y-axis of the 1/v_0_ versus 1/[S] plot. Besides that, the slope of those lines (*K*_s_ / *k*_3 app_) increase up to a limiting value, 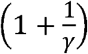,at infinite [ I], producing an hyperbolic curve in the *K*_s_/*k*_3 app_ versus [I] plot (Supplementary Figure 3). These are exactly the same features observed for a partial competitive inhibitor, as observed for Imidazole in the presence of the wild-type Sfβgly (Figure 1) [3].

Furthermore, when the model presented in Figure 10 is interpreted by assuming β = 0, that indicates an ESI complex totally unproductive, the general equation 1 reduces to the form presented in the equation 3 (Supplementary Figure 3):

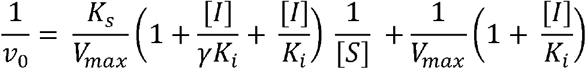

Thus, equation 3 shows that if the ESI complex does not form product, the lines in the Lineweaver-Burk plot crosses at the second quadrant, whereas their slopes (*K*_s_/*k*_3 app_) and intercepts (1/*k*_3 app_) show a linear growth in the presence of increasing [I] (Supplementary Figure 3). These are the properties observed for a linear mixed inhibitor, as Tris is for the wild-type Sfβgly (Figure 1) [1].

Therefore, the previously detected inhibition mechanisms for Imidazole and Tris, partial competitive and linear mixed, respectively, traditionally understood as single binding site mechanism can indeed arise from the same inhibitor interaction within two different sites in the wild-type Sfβgly following the premises expressed in the general model here presented (Figure 10).

### The general model explains the Imidazole and Tris inhibition of the mutant Sfβgly

Interestingly, recalling that the scheme presented in the Figure 10 was designed as a general model, one could then assume a condition not discussed yet, i.e. 0 < β <1. In this situation, in which the ESI complex is less productive than the ES, the general equation 1 is applied without modifications. In that format, equation 1 indicates that lines in the Lineweaver-Burk plot crosses in the second quadrant, whereas their slopes (*K*_s_/*k*_3 app_) and intercepts (1/*k*_3 app_) show a hyperbolic growth in the presence of increasing [I], reaching plateaus at infinite [I], 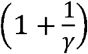 and 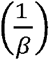,respectively (Supplementary Figure 3). Those are the features of a partial mixed inhibitor, as observed for the F251A mutant when using Imidazole as inhibitor (Table 1; Figure 5). Hence the effect of the F251A replacement was a modification of the β factor from 1 to 0.6 (Table 1), which in the context of the general equation 1 is enough to generate a partial mixed mechanism from a partial competitive one, previously seen for the wild-type (Table 1; Figures 1 and 5).

Similar discussion can be elaborated to the inhibition of the S247Y mutant by Imidazole (Figure 2; Table 1). As mentioned above, when the β factor drops to 0, this modification converts the general equation 1 from a partial competitive (β = 1) to linear mixed (β = 0; Equation 3). So, the effect of the mutation S247Y seems to be the inactivation of the ESI complex involving Imidazole.

Thus, the general model (Figure 10 and Equation 1) also applies to understand the mutational effects on the inhibition mechanisms. Particularly as presented above, different mechanism, partial competitive, partial mixed and linear mixed, interconvert simply by modifications in the reactivity of the ESI complex, expressed in the β factor that multiplies *k*_3_. So, mutations F251A and S247Y may have changed the Imidazole interactions and positioning in the lateral pocket of the Sfβgly active site, which triggered an alteration in the nearby interactions of the substrate transition state within the active site, consequently decreasing the stability of the ESI complex that was manifested in *k*_3_ variations and expressed by the β factor. That agrees with the understanding that enzymes preferentially bind the transition state of the substrate [7].

Now to complete the analysis of how the general model explains the emergence of the different inhibition mechanisms upon mutation, it still remains to be approached the N249A and N249Q mutants in the presence of Imidazole and S247Y, N249A and N249Q in the presence of Tris (Table 1 and 2). In these cases, a linear competitive mechanism was systematically observed.

A qualitative inspection of the general model (Figure 10) suggests that if an inhibitor binds more strongly within the active site, forming the IE complex, and the inhibitor concentration is lower than that dissociation constant of the second binding site, the relative concentration of the remaining complexes, EI and ESI, will be insignificant. This situation can be quantitatively approached by setting γ << 1 and β = 1 when [I] < *K*_i_ in the general equation 1, which is simplified to equation 4 (Supplementary Figure 3):

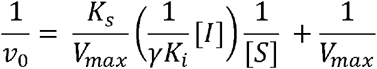

Such combination, γ << 1 and β = 1, suggests that the mutations N249A and N249Q on the Imidazole inhibition (Table 1) and also S247Y, N249A and N249Q on the Tris (Table 2), basically disrupted Imidazole and Tris interactions around the active site opening, making the binding in the interior of the active site relatively much more favorable.

Finally, recalling again that the scheme presented in the Figure 10 was designed as a general model, one could then analyze combinations of β and γ factors not discussed yet. For instance, assuming β = 0 and γ >> 1, which means an unproductive ESI complex and higher inhibitor affinity for the binding site around the active site opening, the equation 3 above simplifies to the equation 5 (Supplementary Figure 3):

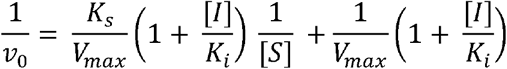

As seen, equation 5 describes a linear non-competitive inhibition mechanism (intersecting, linear, non-competitive; [1]), characterized by lines in the Lineweaver-Burk plot with (*K*_s_/*k*_3 app_) and (1/*k*_3 app_) showing a linear growth in the presence of increasing [I] (Supplementary Figure 3). Besides that, lines observed at different [I] crosses at the same point in the 1/[S] axis, as shown by the same term multiplying both the slope and intercept in the equation 5.

A final combination is 0 < β < 1 and γ >> 1, which means an ESI less productive than the ES complex combined to a higher inhibitor affinity for the binding site around the active site opening. In that condition, the equation 1 transforms into the equation 6 (Supplementary Figure 3):

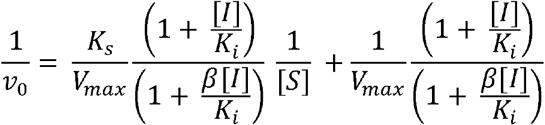

The equation 6 describes lines in a Lineweaver-Burk plot that present increasing slopes and 1/ ν_0_ intercepts as the [I] rises. In addition, such increment is hyperbolic and at infinite [I] both parameters reach a limiting plateau equal 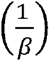 to (Supplementary Figure 3). Those are features of partial non-competitive inhibitors (intersecting, hyperbolic, non-competitive; [1]).

Then considering these two final situations and those discussed above for the mutant and wild-type Sfβgly, the foundations of the general model (Figure 10) engendered six different inhibition mechanisms (3 linear: competitive, non-competitive and mixed and 3 hyperbolic or partial: competitive, non-competitive and mixed).

### The γ factor explains the apparent hindrances between I and S in the formation of the ESI complex

Notably, the presence of the γ factor confers interesting features to the general model (Figure 10). For instance, considering the comparison of two situations in which γ < 1 and γ = 1, in the former a larger part of the enzyme population is diverted to the IE complex. Thus, in the former, lower concentrations of the ESI and ES complexes are observed in the same [S] and [I]. Consequently, the presence of the IE complex introduces an effect of apparent *K*_i_ and *K*_s_ increments because higher [I] and [S] will be needed to reach the same levels observed for the situation in which γ = 1. Traditionally such apparent *K*_i_ and *K*_s_ increments observed in inhibition mechanisms that involve an ESI complex are interpreted as a mutual hindrance between S and I in the formation of the ESI complex. To account for that, an α factor (α > 1) is usually introduced multiplying the *K*_i_ and *K*_s_ in those kinetic models and equations [1].

Nevertheless, here we propose that the mutual hindrance between S and I is just an apparent effect due to the carriage of part of the enzyme population to the IE complex, which goes back to a clear affinity difference between the two binding sites of the inhibitor (γ < 1). Consequently, the traditional α factor can be expressed in terms of the γ factor (Equation 5; Supplementary Figure 3).

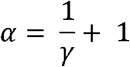

This inverse relation indicates that as the inhibitor affinity for the active site becomes dominant, which is represented as the γ factor decrease, the α factor gets higher than 1. Hence, an apparent mutual hindrance between S and I is observed (Supplementary Figure 3). Oppositely, as the inhibitor preferentially binds elsewhere beyond the active site, the γ factor increases and as a consequence α tends to 1. So, the apparent hindrance disappears.

In brief, no physical obstruction between S and I occurs in the formation of ESI complexes, in fact the apparent hindrance between S and I emerges from the inhibitor relative affinity for two independent binding sites.

Based on that, Tables 1 and 2 can be red again by converting the α factor in the relative binding affinity of the Imidazole and Tris within and outside the active site, as expressed in the γ factor (Table 3).

**Table 3.** Relative binding affinity of inhibitors within and outside the active site of Sfβgly.

| inhibitor | enzyme | γfactor |
| --- | --- | --- |
| Imidazole | wild-type | 0.5 |
|  | S247Y | 0.4 |
|  | F251A | 0.5 |
| Tris | wild-type* | 0.5 |
|  | F251A | 0.09 |

The γ factor < 1 indicates that the inhibitor has higher binding affinity for the active site in comparison to the interaction around the active site opening. The opposite holds for γ factor < 1.

In brief, Table 3 indicates that the Imidazole and Tris have a two-fold higher affinity for the active site of the wild-type Sfβgly when compared to the binding around the active site opening, as in the lateral pocket. Indeed, based on equations 1 and 2 (above) by isolating the terms multiplying the *K*_s_/*V*_max_ and converting them into lines (Supplementary Figure 4), the dissociation constants for the inhibitor bound into the active site (γ*K*_i_) and around its opening (*K*_i_) were calculated as 22.5 and 45 mM for imidazole and 6 and 12 mM for Tris, respectively. These determinations indicating that those inhibitors have more affinity for the active site agrees with the observation of Tris within the Sfβgly active site in crystallographic structures [2].

## Conclusions

The general model depicted in the Figure 10 and Equation 1 is robust enough to be the foundation for emergence of the complete set of inhibition mechanisms shown in Table 1 and Table 2 and Supplementary Figure 1. The two interaction sites for the inhibitor featuring in the model agree with the structural data (crystallographic and computational) showing Tris and Imidazole binding in two different regions: within the active site and around the active site opening [3; 4]. Moreovere, the relative affinity of Tris for these two sites calculated the general model equation shows a preferential binding in the active site, which agrees with the crystallographic data [2]. Modifications of the β factor generate partial competitive, partial mixed and linear mixed inhibition mechanisms. The β factor changes merely reflect differences in the ESI reactivity, which is in line with the perspective that enzymes preferentially bind the substrate transition state [7]. Besides that, combined modifications of the γ and β factors generate linear competitive, linear non-competitive and partial non-competitive inhibition mechanism.

In brief, the general model does not only explain the inhibition data presented here for the wild-type and mutant Sfβgly, it indeed bring together on top of the same theoretical foundation six fundamental inhibition mechanisms previously described in the literature.

The statement that small molecule inhibitors may found more than one binding site in an enzyme structure is realistic. A range of relative affinities for those sites, represented in the γ factor, is a natural consequence. The assumption that inhibitor and enzyme interactions may be manifested in a range of effects on the enzyme interactions with the substrate transition state, which is encapsulated in the β factor, is also reasonable. Therefore, the general model (Figure 10; Equation 1) is a realistic perspective of the small molecule inhibitors interaction with enzymes, explaining a broad range of phenomena previously attributed to diverse mechanisms.

In short, we conclude that six fundamental inhibition mechanisms, linear and hyperbolic, are facets of the same general model here presented.

## 2. Material and Methods

### 2.1 Site-directed Mutagenesis Experiments

Site-directed mutagenesis experiments were performed using QuikChange Lightning Site-Directed Mutagenesis Kit (Agilent Technologies, Santa Clara, CA, USA) according to manufacturer instructions. Mutations were confirmed by DNA sequencing.

### 2.2 Expression and Purification

The Sfβgly mutants were produced and purified as previously reported [3]. The purified mutant Sfβgly samples were subjected to buffer exchange using PD Minitrap G-25 columns (Cytiva, Marlborough, MA, USA). The final samples were stored in 100 mM phosphate buffer pH 6 at 4°C. Sample homogeneity was assessed by SDS-PAGE [8] and protein concentration was measured using the bicinchoninic acid (BCA) assay [9].

### 2.3 Kinetic Assays

#### 2.3.1 Inhibition of Sfβgly mutants by Imidazole

The initial rate of hydrolysis (ν_0_) of at least 10 different concentrations of p-nitrophenyl β-glucoside (NPβglc), ranging from 0.625 to 25 mM, was determined in the presence of 5 different Imidazole concentrations (0 to 120 mM).

Initial rates were determined from three separate reactions with a single enzyme sample performed at 30°C. NPβglc and Imidazole were prepared in 100 mM phosphate buffer, pH 6.0. The product, *p*-nitrophenolate, was detected as previously described [3]. NaCl was added to adjust the ionic strength among reactions, with 120 mM Imidazole as the reference. Data were analyzed using Lineweaver–Burk plots. Linear fits were accepted when showing *R*^2^ values higher than 0.9. The *K*_i_ and α factor were determined using procedures appropriated for the inhibition mechanism [1] and are expressed as median and standard deviation.

#### 2.3.2 Inhibition of Sfβgly mutants by Tris

This section describes experiments similar to those in 2.3.1, with the only difference being the use of Tris instead of Imidazole as the inhibitor. The initial rates of hydrolysis of several p-nitrophenyl β-glucoside (NPβglc) concentrations were determined at various concentrations of Tris (0 to 120 mM) under the same conditions and procedures.

### 2.4 Molecular Docking

The crystallographic structure of the Sfβgly (PDB 5CG0) was used in the molecular docking [2] as previously described [3; 4]. The coordinates for Imidazole and Tris were generated using Gabedit v.2.5.1 [10], and both were docked in their mono-protonated states (+1 charge).

## Abbreviations

Sfβgly: GH1 β-glucosidase from *Spodoptera frugiperda* (PDB 5CG0)
NPβglc: *p*-nitrophenyl β-glucoside
Tris: 2-amino-2-(hydroxymethyl)-1,3-propanediol

## Funding sources

This project was supported by FAPESP (Fundação de Amparo à Pesquisa do Estado de São Paulo; Grants 2021/03967-6 and 2021/10577-0), CAPES (Coordenação de Aperfeiçoamento de Pessoal de Nível Superior) and CNPq (Conselho Nacional de Desenvolvimento Científico). The funders had no role in study design, data collection and analysis, decision to publish, or preparation of the manuscript.

## Author Contributions

**Rafael S. Chagas:** conceptualization; formal analysis; investigation; writing original draft; writing review and editing. **Sandro R. Marana:** conceptualization; funding acquisition; resources; formal analysis; investigation; writing -original draft; writing - review and editing.

## Competing interest

Authors declare no conflicts of interest.

## Supplementary Material

**Supplementary Figure 1.**
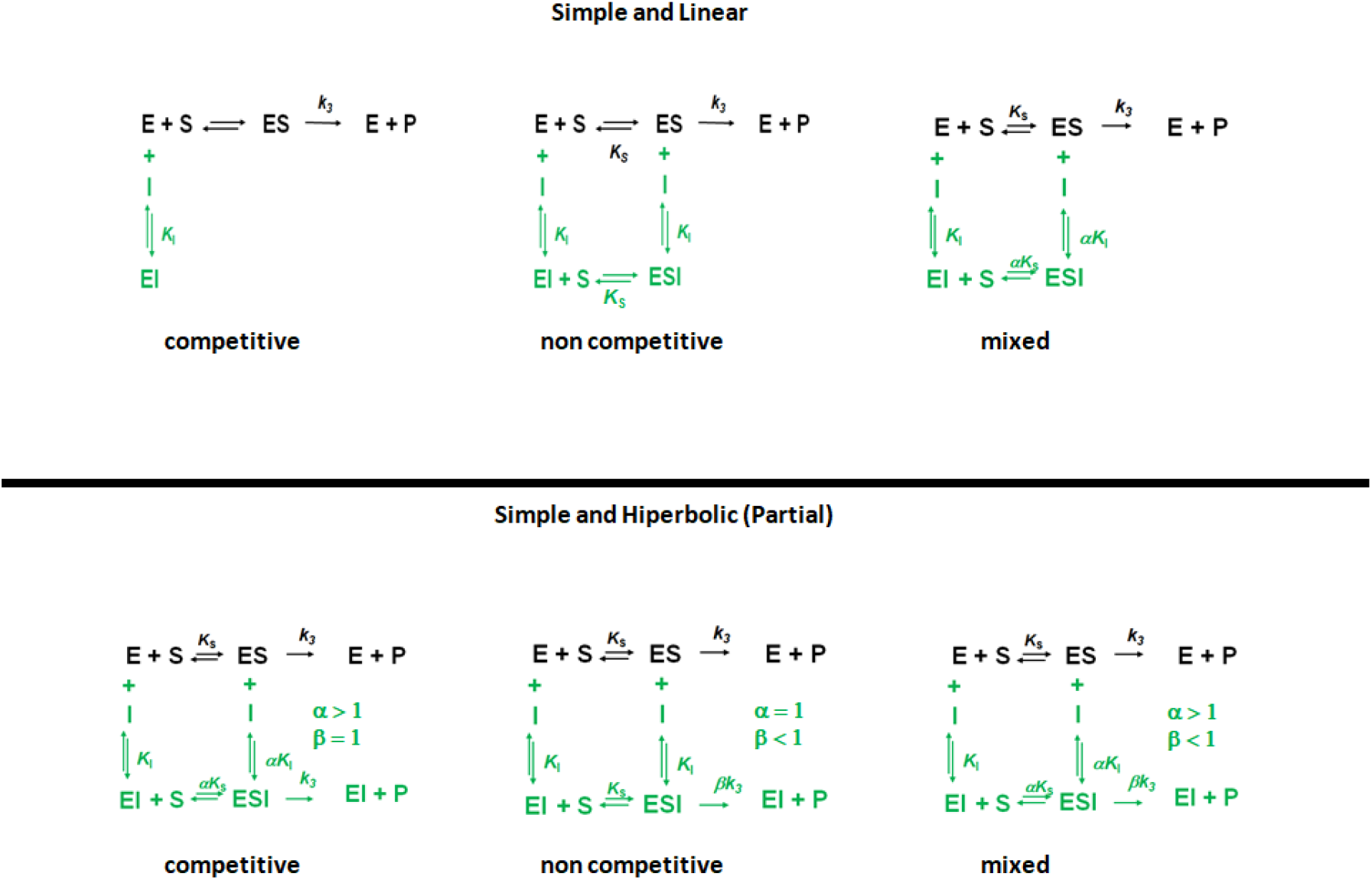
Simple and intercepting inhibition mechanisms

**Supplementary Figure 2.**
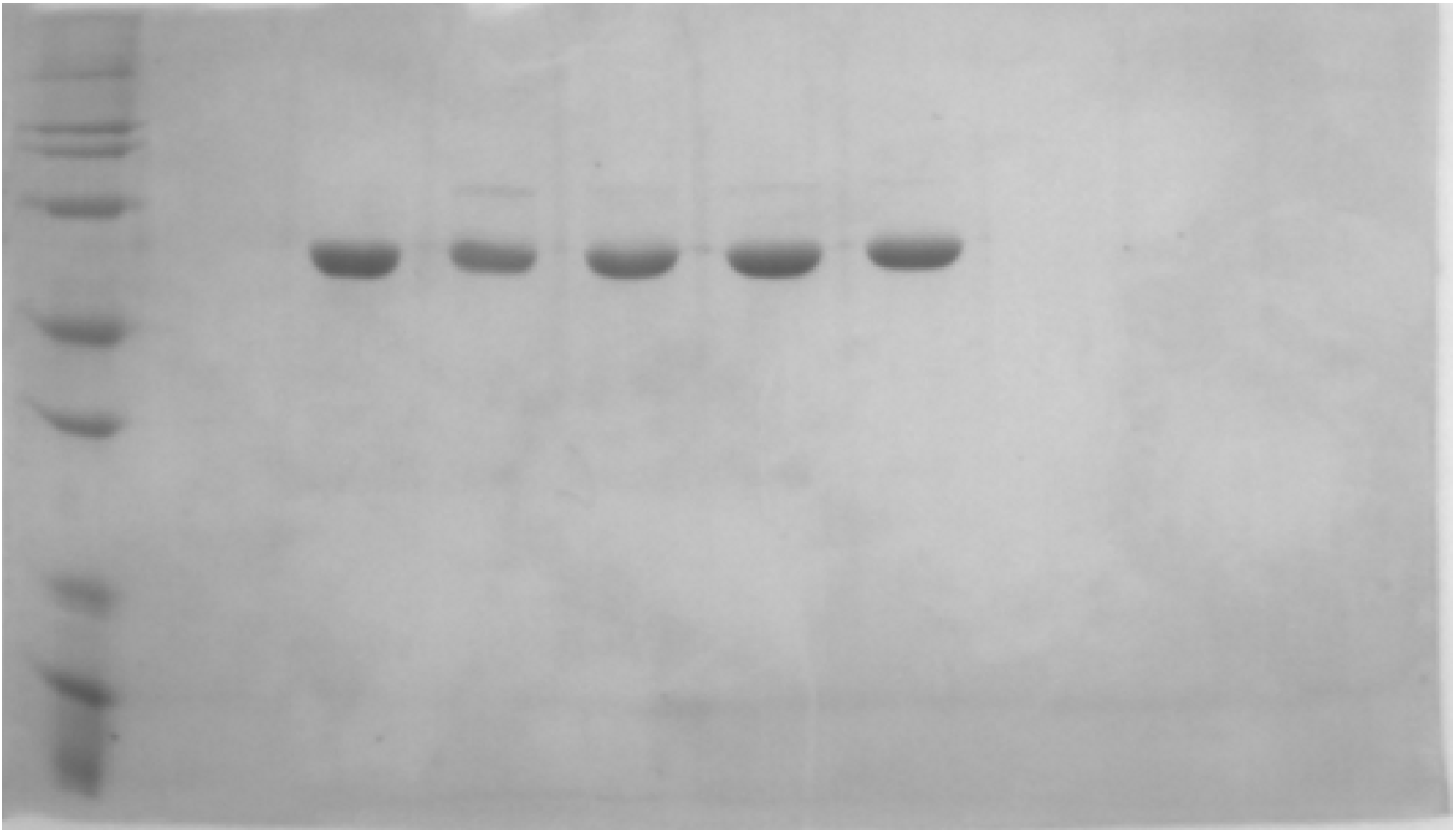
SDS-PAGE of the purified recombinant Sfβgly mutants. Lanes: M – Molecular weight marker (kDa). Sfβgly mutants are: 1 – S247T; 2 – S247Y; 3 – N249A; 4 – N249Q and 5 – F251A. Mutant S247T was not studied in this manuscript.

**Supplementary Figure 3.**
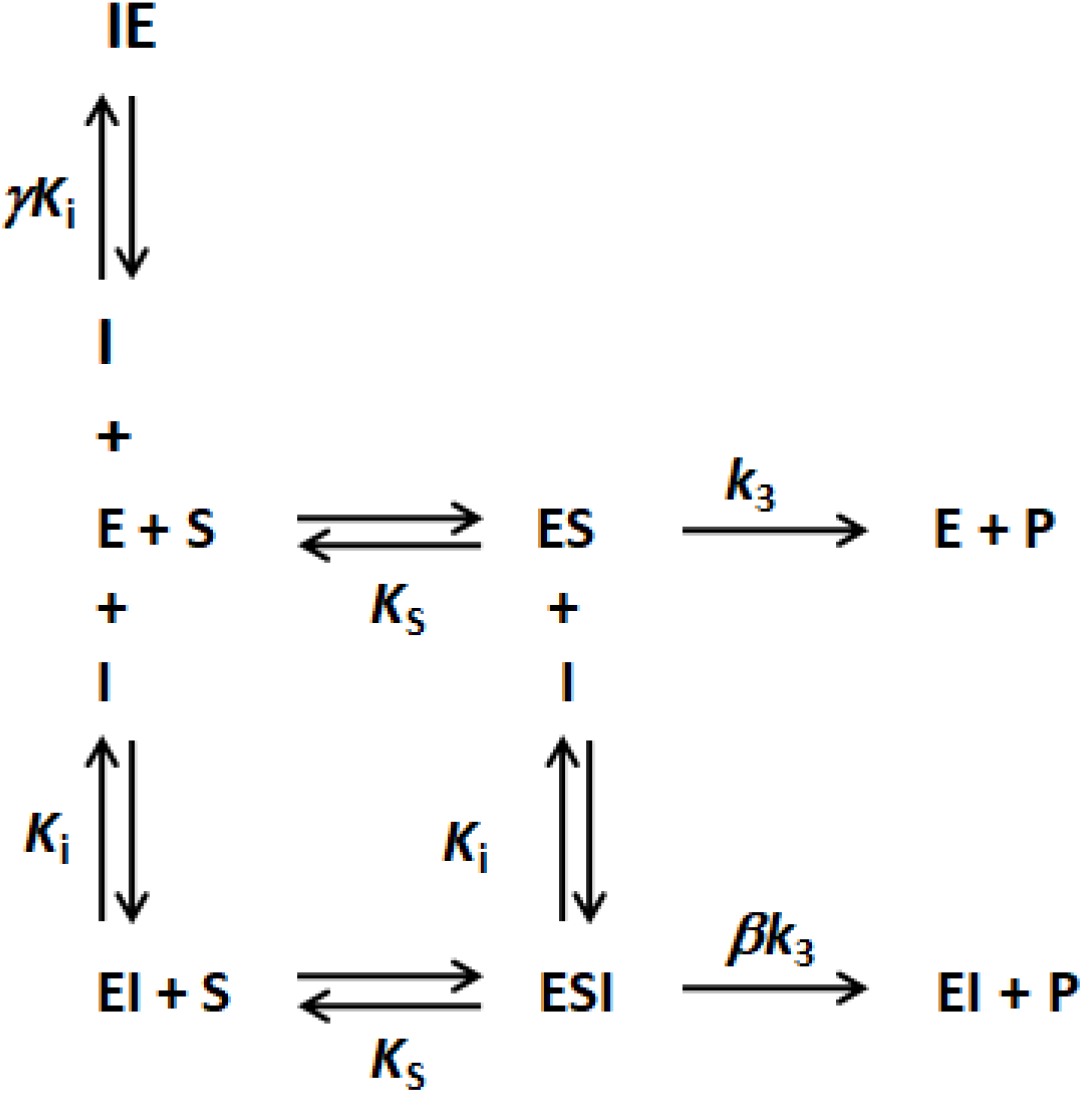
General Mechanism of Enzyme Inhibition: Equations and Deductions. Inhibition mechanism based on two different binding sites of the inhibitor in the enzyme. One of the sites is within the active site, where the inhibitor interaction blocks the substrate binding and results in the IE complex. The second site is beyond the active site, where the inhibitor binding does not affect the substrate interaction within the active site. This binding produces the EI complex. E, free enzyme; ES, enzyme-substrate complex; IE, complex presenting the inhibitor bound within the enzyme active site; EI, complex in which the inhibitor is interacting beyond the active site; ESI, ternary complex presenting the substrate in the enzyme active site and the inhibitor bound elsewhere; γ factor represents the inhibitor relative affinity for the two binding sites available in the enzyme; β represents the inhibitor effect on the *k*_3_.

Based on this scheme, the dissociation constants and the respective complexes are defined as seem below.

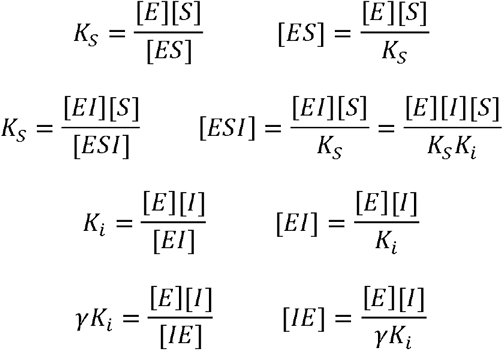

The initial (ν_0_) and maximum rate (*V*_max_) are expressed as:

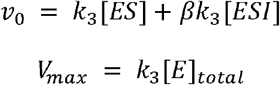

The total enzyme population ([E]_total_) is distributed among the species presented in the scheme above.

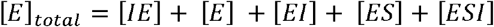

Based on these definitions the relative rate ν_0_/*V*_max_ is expressed as:

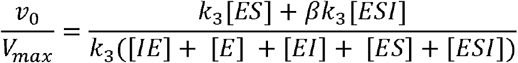

Definitions of the complexes ES, ESI, EI, IE e ESI are inserted into the ν_0_/*V*_max_ equation.

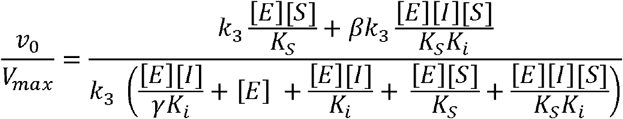

The equation above is simplified by eliminating *k*_3_ and [E]

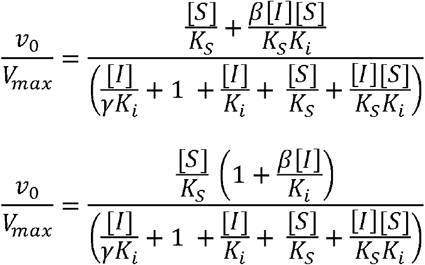

Numerator and denominator are multiplied by *K*_s_.

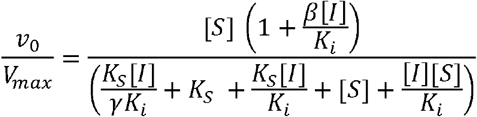

In the denominator, find a common factor in the terms containing *K*_s_ and [S].

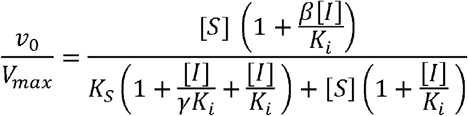

Numerator and denominator are divided by 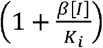.

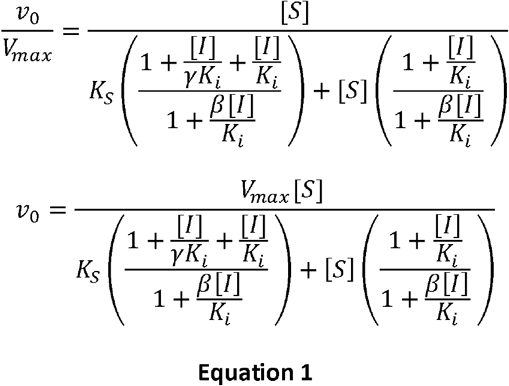

The equation 1 is similar to the Michaelis-Menten equation. Hence, it is linearized using the Lineweaver-Burk inversion.

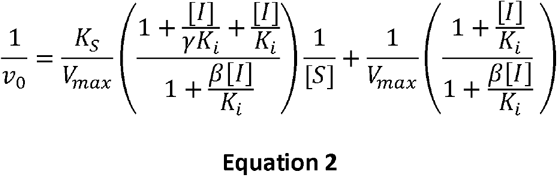

### Case 1 – General Model in β = 1

Taking the general model above and assuming β = 1, the dissociation constants and the respective complexes are defined as:

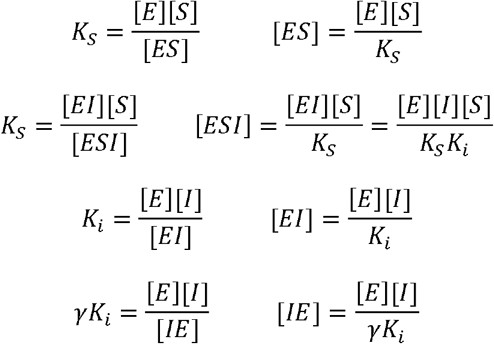

The initial (ν_0_) and maximum rate (*V*_max_) are expressed as:

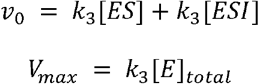

The total enzyme population ([E]_total_) is distributed among the species presented in the scheme above.

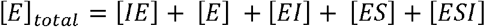

Based on these definitions the relative rate ν_0_/*V*_max_ is expressed as:

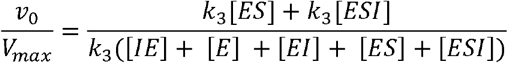

Definitions of the complexes ES, ESI, EI, IE e ESI are inserted into the *ν*_0_/*V*_max_ equation.

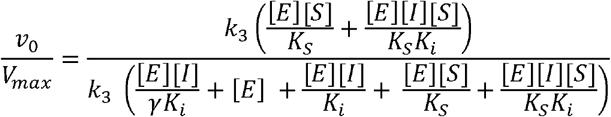

The equation above is simplified by eliminating *k*_3_ and [E]

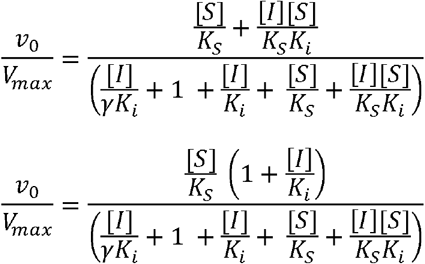

Numerator and denominator are multiplied by *K*_s_.

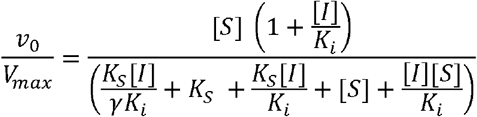

In the denominator, find a common factor in the terms containing *K*_s_ and [S].

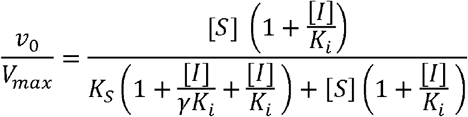

Numerator and denominator are divided by 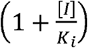.

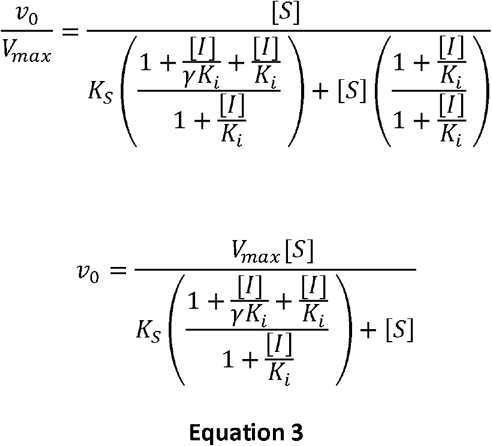

The equation 3 is similar to the Michaelis-Menten equation. Hence, it is linearized using the Lineweaver-Burk inversion.

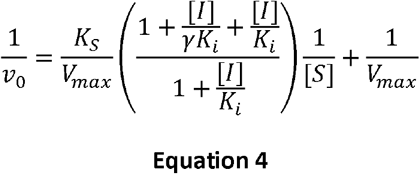

Based on the equation 4, by isolating the term that multiplies the slope (*K*_s_/*V*_max_), we can analyze the behavior of this term as a function of [I].

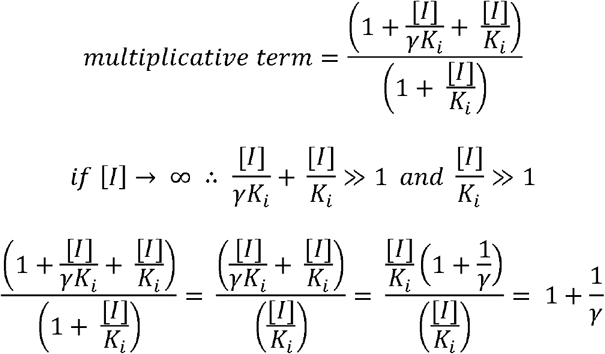

Thus, the multiplicative term has a finite and constant value at infinite [I].

In addition to that, we can simulate the behavior of this term as a function of [I].

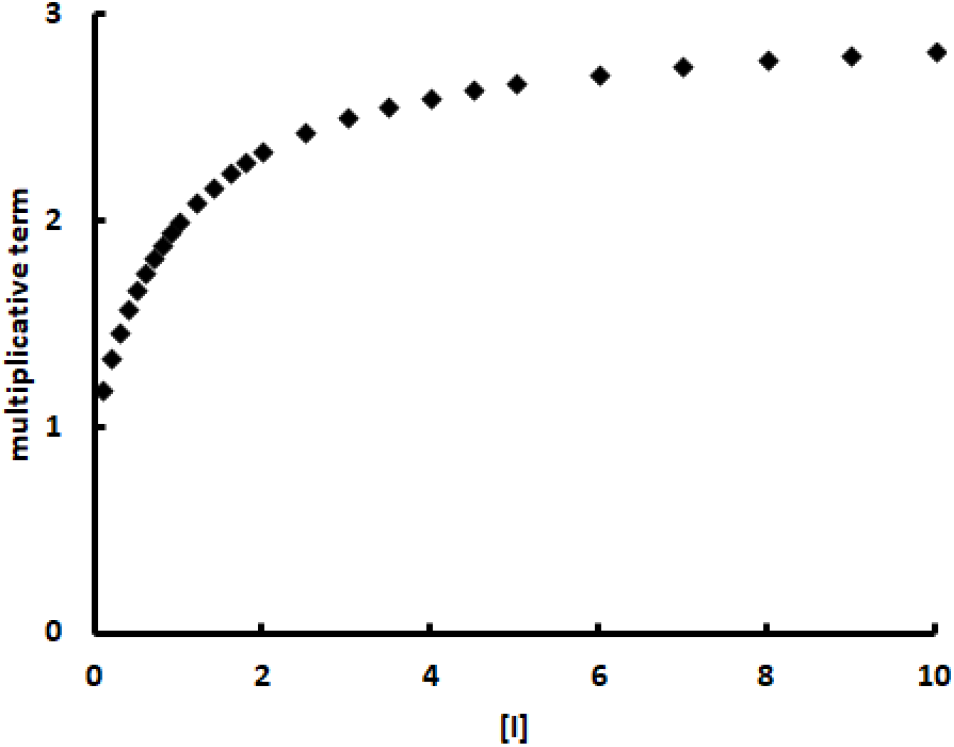

Simulation of the multiplicative term magnitude as a function of [I]. *K*_i_ = 1 and γ = 0.5

### Case 2 – General Model in β = 0

Taking the general model above and assuming β = 0, the dissociation constants and the respective complexes are defined as:

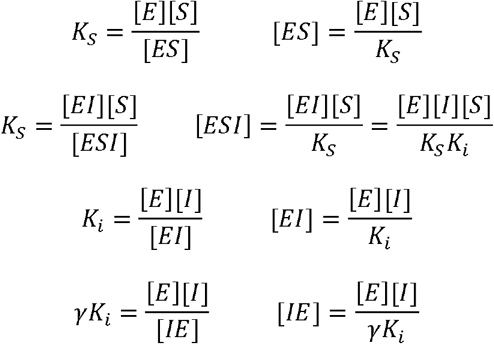

The initial (ν_0_) and maximum rate (*V*_max_) are expressed as:

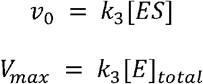

The total enzyme population ([E]_total_) is distributed among the species presented in the scheme above.

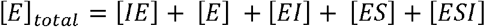

Based on these definitions the relative rateν_0_/*V*_max_ is expressed as:

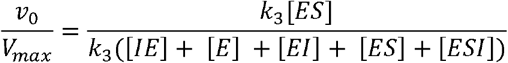

Definitions of the complexes ES, ESI, EI, IE e ESI are inserted into the ν_0_/*V*_max_ equation.

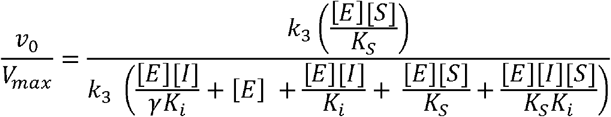

The equation above is simplified by eliminating *k*_3_ and [E]

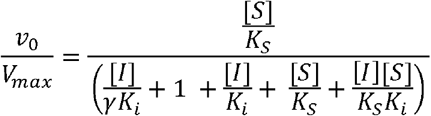

Numerator and denominator are multiplied by *K*_s_.

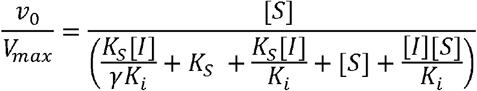

In the denominator, find a common factor in the terms containing *K*_s_ and [S].

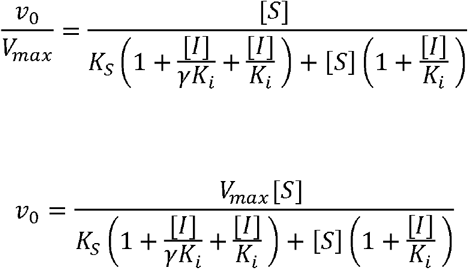

The equation above is similar to the Michaelis-Menten equation. Hence, it is linearized using the Lineweaver-Burk inversion.

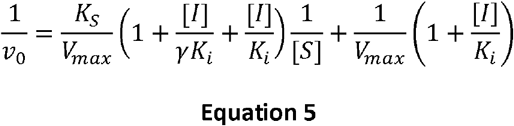

Based on the equation 5, by isolating the terms that multiply the slope (*K*_s_/*V*_max_) and the intercept (1/*V*_max_), we can analyze the behavior of these terms as a function of [I].
Slope multiplicative term

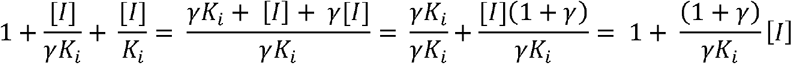

Intercept multiplicative term

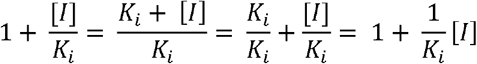

Both terms are linear functions of [I]

### Case 3 – General Model in 0 < β < 1

Taking the general model above and assuming 0 < β < 1, the general equation 1 applies.

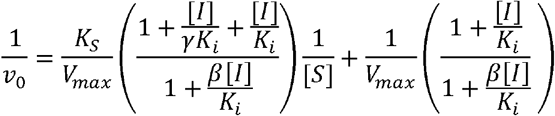

As already demonstrated above, by isolating the term that multiplies the slope (*K*_s_/*V*_max_), we can analyze its behavior as a function of [I].

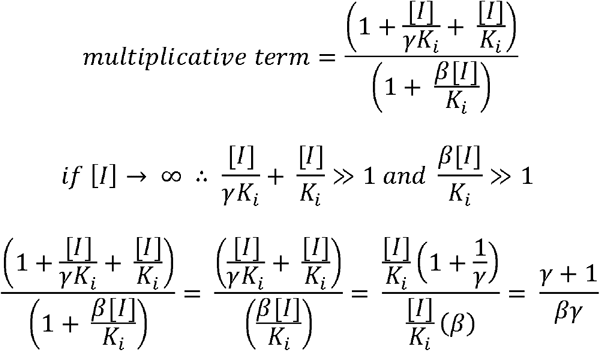

Thus, the slope and intercept multiplicative terms have a finite and constant value at infinite [I]. In addition to that, we can simulate the behavior of this term as a function of [I].

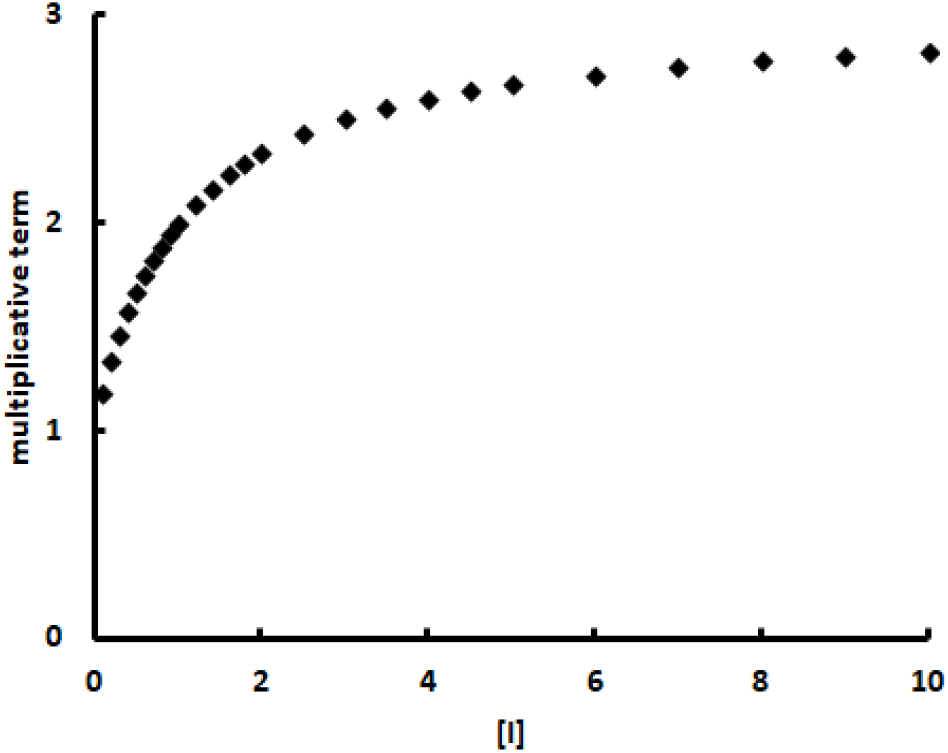

Simulation of the slope multiplicative magnitude as a function of [I]. *K*_i_ = 1 and γ = 0.5

Similarly, by isolating the term that multiplies the intercept (1/*V*_max_), we can analyze its behavior as a function of [I].

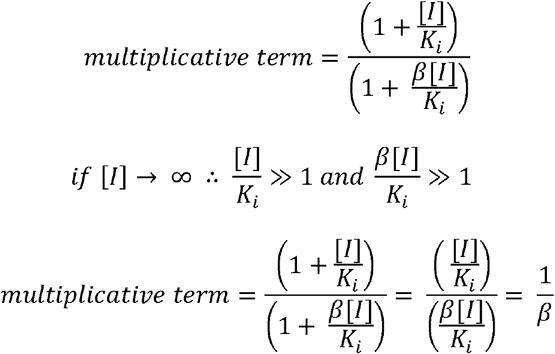

Thus, the intercept multiplicative term has a finite and constant value at infinite [I].

In addition to that, we can simulate the behavior of this term as a function of [I].

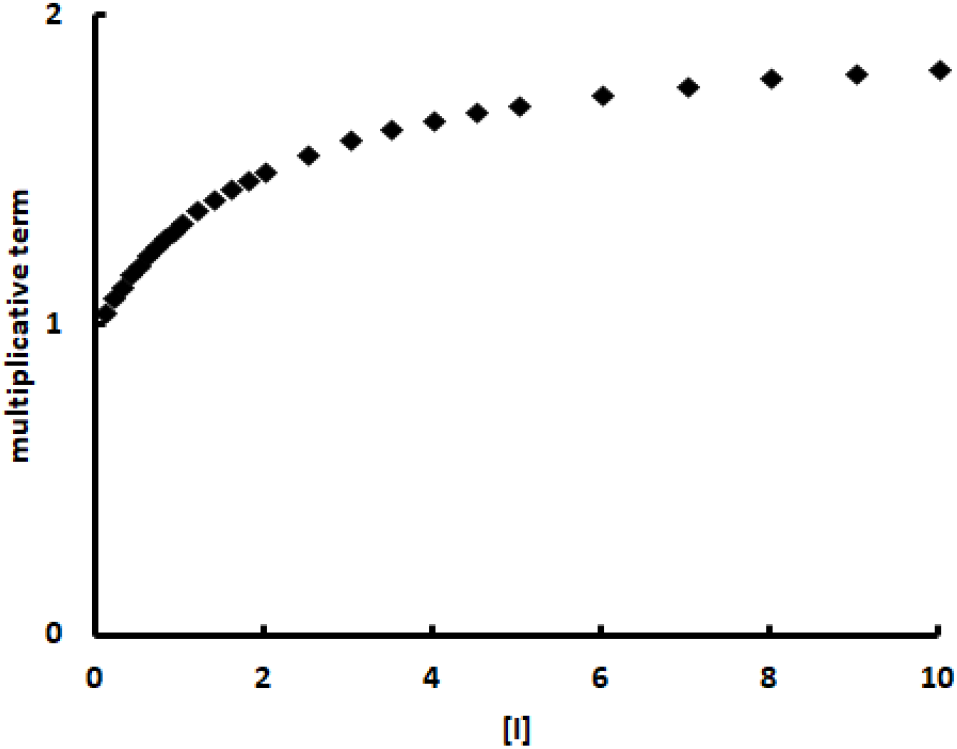

Simulation of the intercept multiplicative magnitude as a function of [I]. *K*_i_ = 1 and β = 0.5

Finally, by assuming arbitrary values for the constants in the Lineweaver-Burk equation 1 above, we can trace the lines observed in two different inhibitor concentrations showing that they meet in the second quadrant.

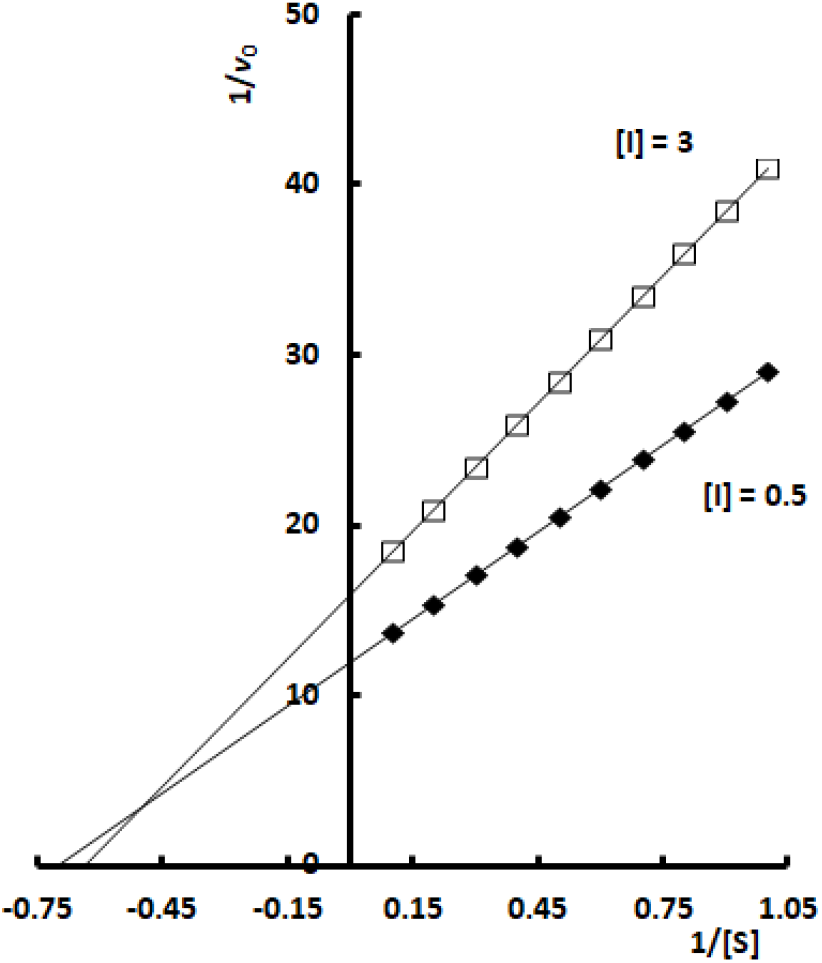

Simulation of the Lineweaver-Burk lines for *K*_s_ = 1, *V*_max_ = 0.1, β = 0, *K*_i_ = 1 and γ = 0.5 in two different [I] (0.5 and 3)

### Case 4 – General Model in β = 1 and γ << 1

Taking the general model and assuming β = 1, the kinetic equation 1 deduced above is

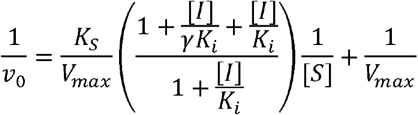

Assuming γ << 1 tem factor multiplying the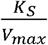 is simplified as follows:

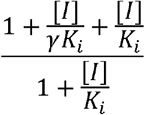

If γ << 1 then 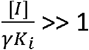 and also 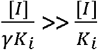. Thus

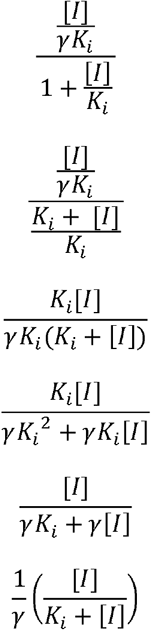

Quando *K*_i_ >> [I], então *K*_i_ + [ I] ≈ *K*_i_

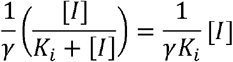

In this particular situation, the kinetic equation is written as

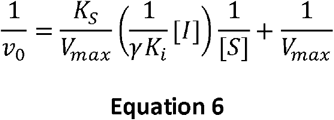

### Case 5 – General Model in β = 0 and γ >> 1

Taking the general model and assuming β = 0, the kinetic equation 1 deduced above is

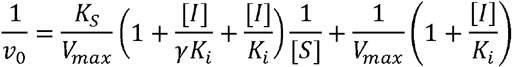

Assuming γ >> 1 the factor multiplying the 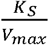 is simplified as follows:

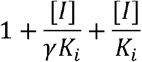

If γ >> 1 then 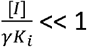,then

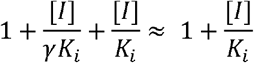

So, the kinetic equation is rewritten as:

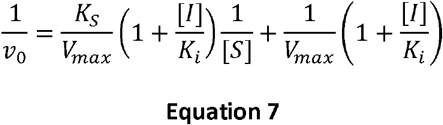

### Case 6 – General Model in 0 < β < 1 and γ >> 1

Taking the general model and assuming 0 < β < 1, the kinetic equation 1 deduced above is

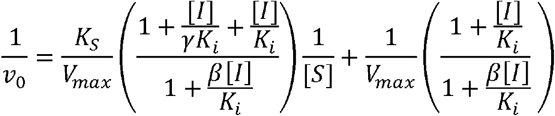

Assuming γ >> 1 the term multiplying the 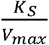 is simplified as follows:

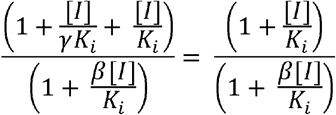

Hence, the slope and intercept are multiplied by the same term. Next we can analyze its behavior as a function of [I].

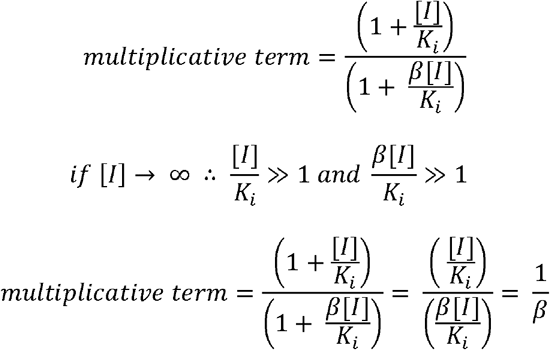

Thus, the intercept multiplicative term has a finite and constant value at infinite [I].

In addition to that, we can simulate the behavior of this term as a function of [I].

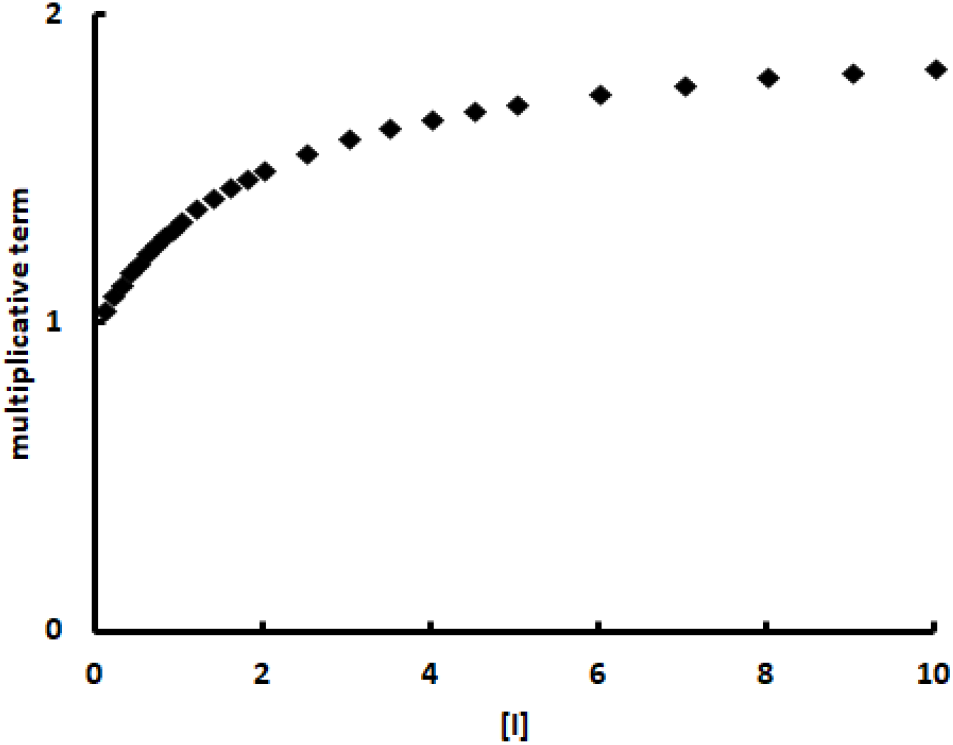

Simulation of the slope and intercept multiplicative terms magnitude as a function of [I]. *K*_i_ = 1 and β = 0.5

### Correlation between parameters γ and α

Assuming β = 1 the general inhibition model generates the equation 4

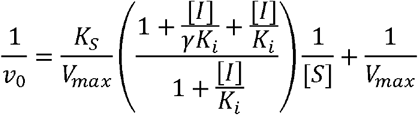

By isolating the term that multiplies the slope (*K*_s_/*V*_max_) and analyzing its behavior as a function of [I], we conclude that at infinite [ I], the term converge to a finite number.

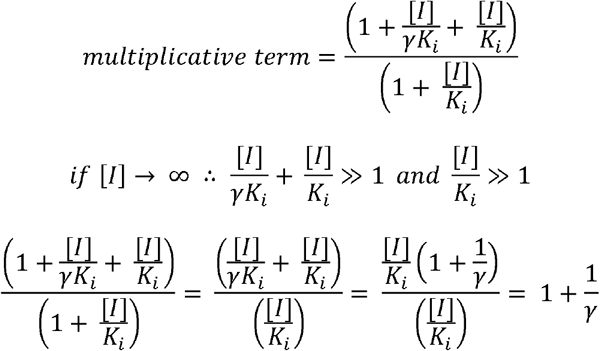

Similarly, the kinetic equation for a partial competitive mechanism has a term that multiplies the slope (*K*_s_/*V*_max_), which also converges to a finite number.

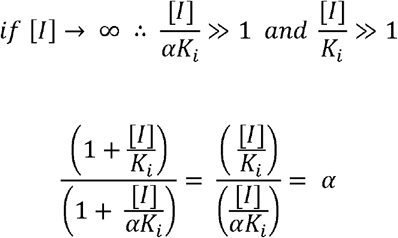

Hence, both multiplicative terms converge to finite numbers, indicating that

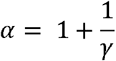

Based on this equation, we may simulate the relation between α and γ

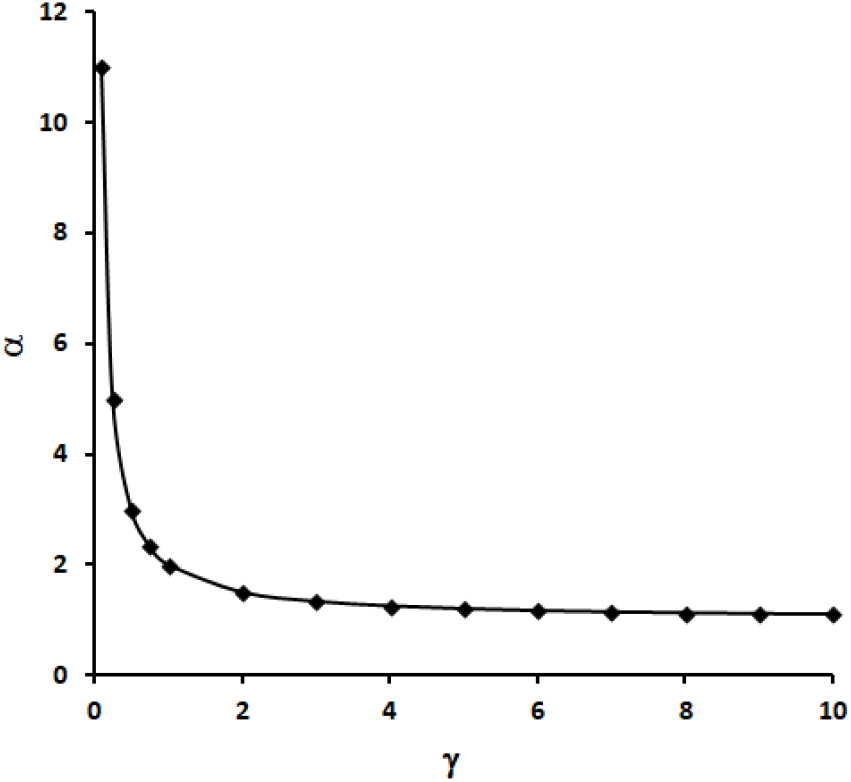

Correlation between parameters γ and α.

### Supplementary Figure 4 – Determination of *K*_i_ using the general model

**Imidazole** is a partial competitive inhibitor of Sfbgly (Table 1). This inhibition mechanism is a facet of the general model when β = 1 (Case 1). Under this condition, the general model results in the equation 4:

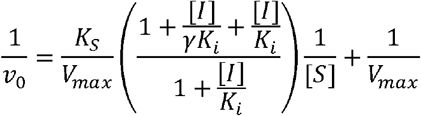

from which the slope is isolated

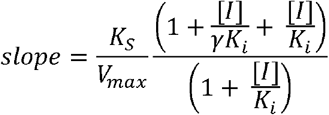

and linearized

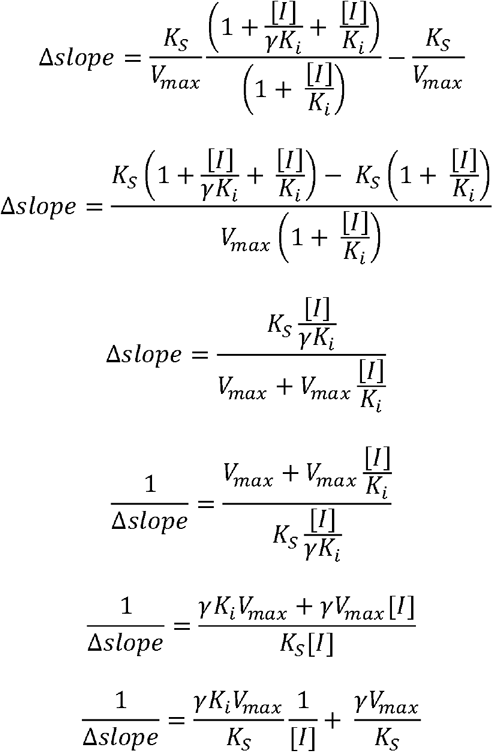

So, if 1/Δslope = 0, then

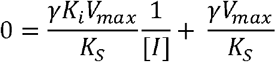

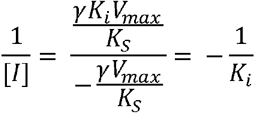

Hence the plot 1/Δslope versus 1/[I] results in a line that crosses the x-axis at -1/*K*_i_. Considering that if β = 1 the general model behaves as a simple partial competitive mechanism, in which the plot 1/Δslope versus 1/[I] results in a line that crosses the x-axis at - 1/α*K*_i_, then

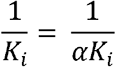

Therefore the α*K*_i_, calculated using the simple partial competitive model (as in Table 1), corresponds to the *K*_i_, if assumed the general model.

**Tris** is a linear mixed-type inhibitor of Sfβgly (Table 1). This inhibition mechanism is a facet of the general model when β = 0 (Case 2). Under this condition, the general model results in the equation 5:

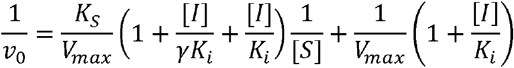

from which the slope is isolated

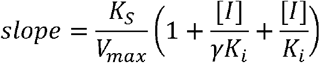

and linearized

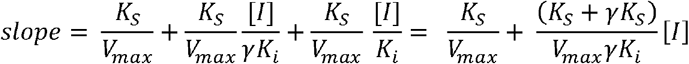

So, if slope = 0, then

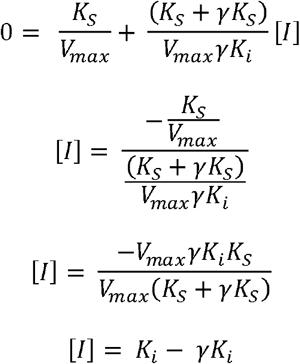

Hence the plot slope versus [I] results in a line that crosses the x-axis at *K*_i_ - γ*K*_i_.

Considering that if β = 0 the general model behaves as a simple linear mixed mechanism, in which the plot slope versus [I] results in a line that crosses the x-axis at K_i_, then

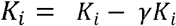

Therefore, the *K*_i_, calculated using the simple linear mixed model (as in Table 1), corresponds to the *K*_i_ - γ*K*_i_, if assumed the general model. Then, taking the *K*_i_ and α presented in Table 1 and the equation

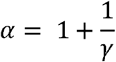

the K_i_ and γ for the general model can be ed.

## Notes

### Competing Interest Statement

The authors have declared no competing interest.

